# Topology identifies concurrent cyclic processes in single-cell transcriptomics and androgen receptor function

**DOI:** 10.1101/2025.01.09.632214

**Authors:** Kelly Maggs, Markus K. Youssef, Cyril Pulver, Jovan Isma, Tâm J. Nguyên, Matthias Arzt, Wouter Karthaus, Heather A. Harrington, Kathryn Hess, G. Paolo Dotto

**Author notes:** Shared first-authorship (Kelly Maggs, Markus K. Youssef). Co-correspondence,.

## Abstract

Standard single-cell RNA-seq analysis frameworks aggregate overlapping biological processes and impose a single parametrization, conflating distinct programs. Here, we introduce a topological framework that detects and disentangles multiple cyclic processes directly from single-cell transcriptomic data. We validate this approach on synthetic datasets and scRNA-seq profiles of human dermal fibroblasts under control conditions and following androgen receptor (AR) silencing, as well as in vivo mouse prostate regeneration under androgen receptor add-back. We show robust cell cycle structure across conditions, identify an unbiased AR-linked stress signature related to the senescence and proliferation across organisms, and uncover cholesterol homeostasis as an AR-linked program in tissue regeneration. This framework enables identification and separation of concurrent cyclic processes from snapshot single-cell data, revealing complex multi-dimensional regulatory dynamics inaccessible to standard clustering analysis.

---

Single-cell RNA-seq gives a high-dimensional point cloud that is often analyzed as an approximation of the Waddington landscape. The long-held promise of single-cell sequencing has been to uncover both ‘intercellular heterogeneity and subtle quantitative processes’ [1]. The standard single-cell analysis pipeline of clustering over low-dimensional projections is remarkably successful at identifying cellular heterogeneity. However, the distillation of subtle biological and regulatory processes remains a subject of intense inquiry and debate [2].

Waddington’s classical epigenetic landscape [3] is a common starting point for mathematical and computational models of development [4–6]. In the landscape model, cells roll down branching valleys carved by underlying regulatory programs. However, such a model lacks a framework for the cyclic processes, such as cell-cycle oscillations, tissue regeneration, and metabolic processes, including stress-response circuits. Additionally, the current model fails to capture that these processes occur concurrently. An updated model would therefore be a sophisticated, high-dimensional mathematical structure built from multiple processes—some one-way (irreversible) and others that are recurrent (cyclic). An unsolved problem in the field is how to identify and separate these different types of processes.

Existing methods in computational single-cell sequencing analysis reflect these conceptual issues in their mathematical modeling assumptions. The vast majority of tools assume that dynamic processes are modeled as 1-dimensional structures such as trees and paths, with some more recent models assuming that the data lies on a circle [7–13]. Projecting onto a single parameter can miss or conflate these concurrent circular programs. While more complicated mathematical models have been proposed in theory [14, 15], the data-driven formalism to describe how multiple processes coalesce to form higher order structure has yet to be laid out in a principled manner.

Here, we rely on recent advances in computational topology to develop a principled mathematical approach to identify distinct cyclic biological processes and study their higher-order interactions. Recent applications of topological data analysis in single-cell studies further support the promise of this topological/geometric perspective [16–21] – particularly the use of cohomology-based circular coordinates in concurrent recent work [22, 23]. Our topological approach relies on persistent cohomology as a mathematical signature of recurrent biological processes. These measurements come equipped with stability theorems that guarantee robustness – a useful feature when working with noisy transcriptomic data. Based on the integration of differential forms and harmonic cocycle [24–27], we introduce new mathematical quantities to measure multi-dimensional recurrent dynamics within the Waddington landscape. We develop a geometric language for analyzing identified topological features to recover pseudo-temporal orderings of both cells and gene expression, allowing an in-depth biological interpretation of the discovered structure.

We first validate that topological and geometric invariants can robustly detect and analyze biological phenomena to yield reliable insight. Using the cell cycle as an example, we introduce a topological-geometric framework to recover the cyclic ordering of cells and dynamic ordering of gene expression consistent with known literature. We show that our predicted ordering is robust across both cell type and organism. Furthermore, this topological approach shows that key cell cycle genes in proliferating human dermal fibroblasts function normally, regardless of whether the androgen receptor is present.

We explicitly demonstrate the existence of multiple concurrent cyclic processes in both synthetic models and real-world transcriptomic data. Specifically, we focus on transcriptional circuits interlocking cell-cycle regulation with androgen receptor (AR)–dependent control of gene expression, where AR functions as a master regulator of developmental and cancer-related transcriptional programs [28]. Topological analysis of androgen modulated prostate regeneration data from [29] reveals that tissue regeneration and cell cycle generate separate circular dimensions Figure 1. The existence of such *multiple recurrent temporal dimensions* challenges the current paradigm of modeling dynamics from single-cell RNA-seq data.

**Figure 1.**
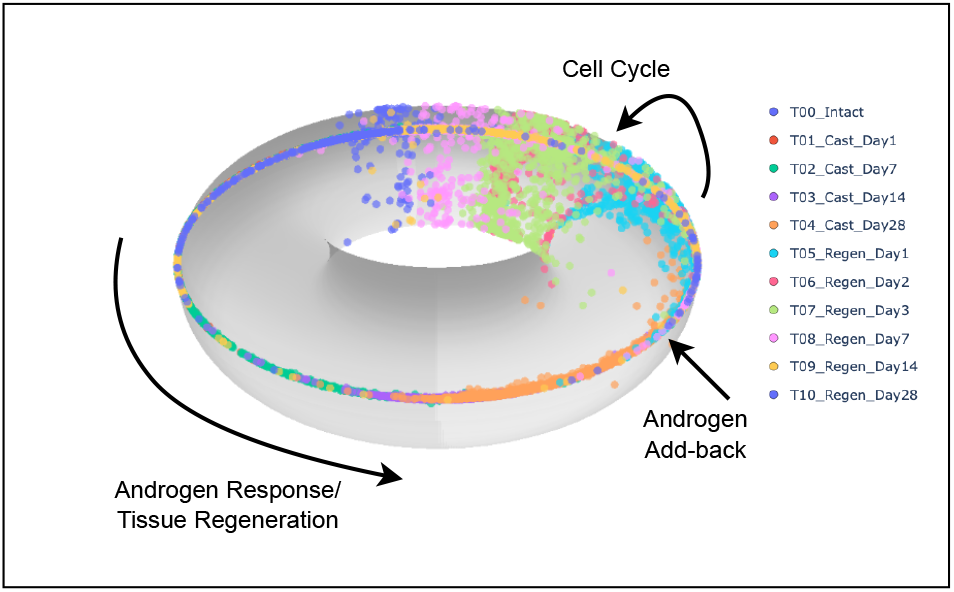
Multi-temporal processes in AR-modulated prostate regeneration. Topological re-analysis of androgen-receptor modulated prostatic regeneration data from [29]. Upon androgen add-back, regeneration and cell cycle span *multiple circular temporal dimensions* in single-cell RNA sequencing data, suggesting *temporal degrees of freedom* in cellular decision making.

Our topological approach shows that key cell-cycle genes in proliferating human dermal fibroblasts remain robustly and coherently regulated regardless of AR status, even though AR acts in these cells as a repressor of senescence-associated, protumorigenic activation [30–32]. We derive an AR stress signature that comprises of upregulation of senescence and pro-inflammatory genes with protumorigenic potential. Cells with AR loss heterogenously exhibit such a signature that is reduced along progression through the cell cycle topology. We further extend our analysis to an in vivo mouse prostate regeneration model under AR control, revealing a consistent interlocking of AR stress signature and topological cell cycle dynamics.

Finally, we apply our cohomology-based circular gene set enrichment to topologically identify cholesterol homeostasis and androgen response as key interlocking sub-processes of tissue regeneration in mouse prostatic epithelial cells. Together, these results establish the broad applicability of our topological framework for identifying and separating complex cyclic biological programs across organisms, tissue contexts, and experimental modalities.

## Results

### Robust topological pipeline for identifying statistically significant circular biological processes

In this work, we introduce *Cocycle Hunter* (`cHunter`) (Figure 2) as a topological framework for single-cell analysis. Circular geometries in high-dimensional single-cell RNA-seq data involve the coordinated behavior of many genes. We extend gene set enrichment, a widely used method in transcriptomic analysis [33], to *circular* gene set enrichment (Figure 2A), which tests whether projecting single-cell data onto a specific gene set produces a statistically significant cyclic pattern. We focus on *gene set projections* by restricting our analysis to groups of biologically relevant genes (e.g., those from the Molecular Signatures Database [34]). This strategy facilitates the detection of recurrent sub-processes and provides a clear basis for downstream biological interpretation.

**Figure 2.**
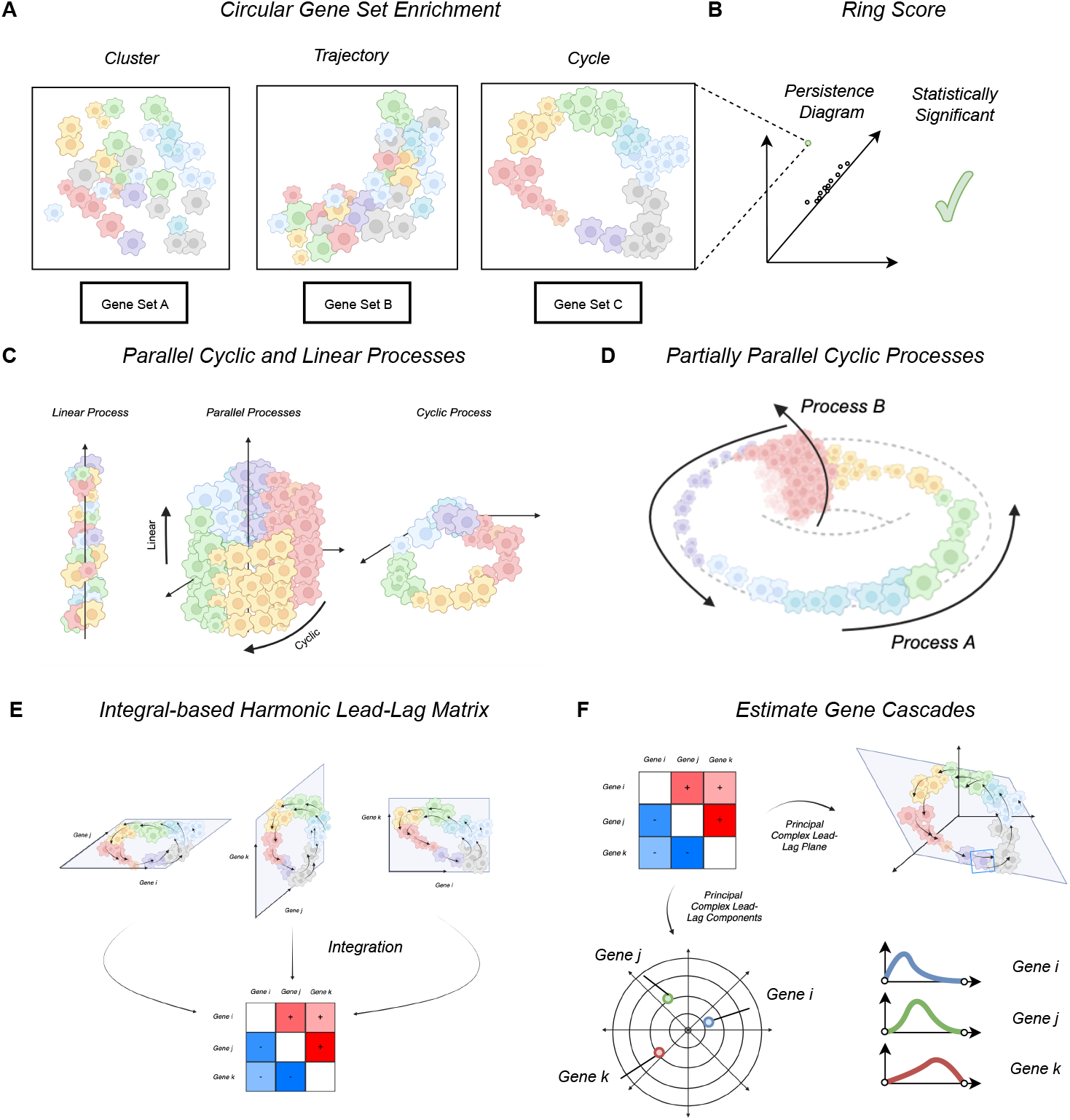
Conceptual overview of cHunter topological and geometric pipeline. **(A)** Circular gene set enrichment tests whether the projection of a single-cell experiment onto a given gene set reveals a cyclic pattern. **(B)** Persistent cohomology is used to quantify circular geometry and statistical significance. **(C)** Topological model of parallel uncoupled linear and circular processes as forming a cylindrical product space. **(D)** Topological model of partially parallel cyclic processes as embedded on the torus. **(E)** The data is further analyzed by projecting onto pairwise gene planes and integrating to compute a lead-lag matrix. **(F)** Finally, eigenvector analysis of this matrix estimates the order of gene activation, revealing gene cascade phases associated with the cycle.

To quantify circularity, we apply a robust mathematical technique, persistent cohomology [35], to automatically detect circular structures within gene set projections and evaluate their significance using a summary statistic, the *ring score* ((Figure 2B). Upon detection of such a significant circular structure, we assign each cell a phase on the inferred cycle, thereby providing an ordering that captures pseudo-dynamics through circular coordinates defining a harmonic signal. These quantities are computed across pairwise gene projections to construct a *lead-lag matrix* (Figure 2F), whose entries encode directional relationships between pairs of genes via integrals describing their orientation in certain subspaces (Figure 2E). Positive and negative values of the *ij*-th entry respectively model expression of gene *i* peaking before or after gene *j* along the circular structure. Spectral analysis of this matrix (Figure 2F) yields an inferred temporal ordering of gene activation, with the amplitude and phase of complex spectral components furnishing quantitative estimates of each gene’s relative contribution and peak activity along the harmonic cycle.

### Topological approach excels at concurrent process analysis

In single-cell RNA-seq data, gene sets may contain more than one dynamic transcriptomic process (Figure 2B). For example, the cell cycle can co-occur with an independent signaling process that induces transcription. In such cases, state-of-the-art methods [8] for estimating circular processes are not designed to disentangle circular dynamics from orthogonal processes. In order to test this hypothesis, we designed a synthetic test dataset (Appendix A.2) consisting of uncoupled circular and linear processes (Figure 3).

**Figure 3.**
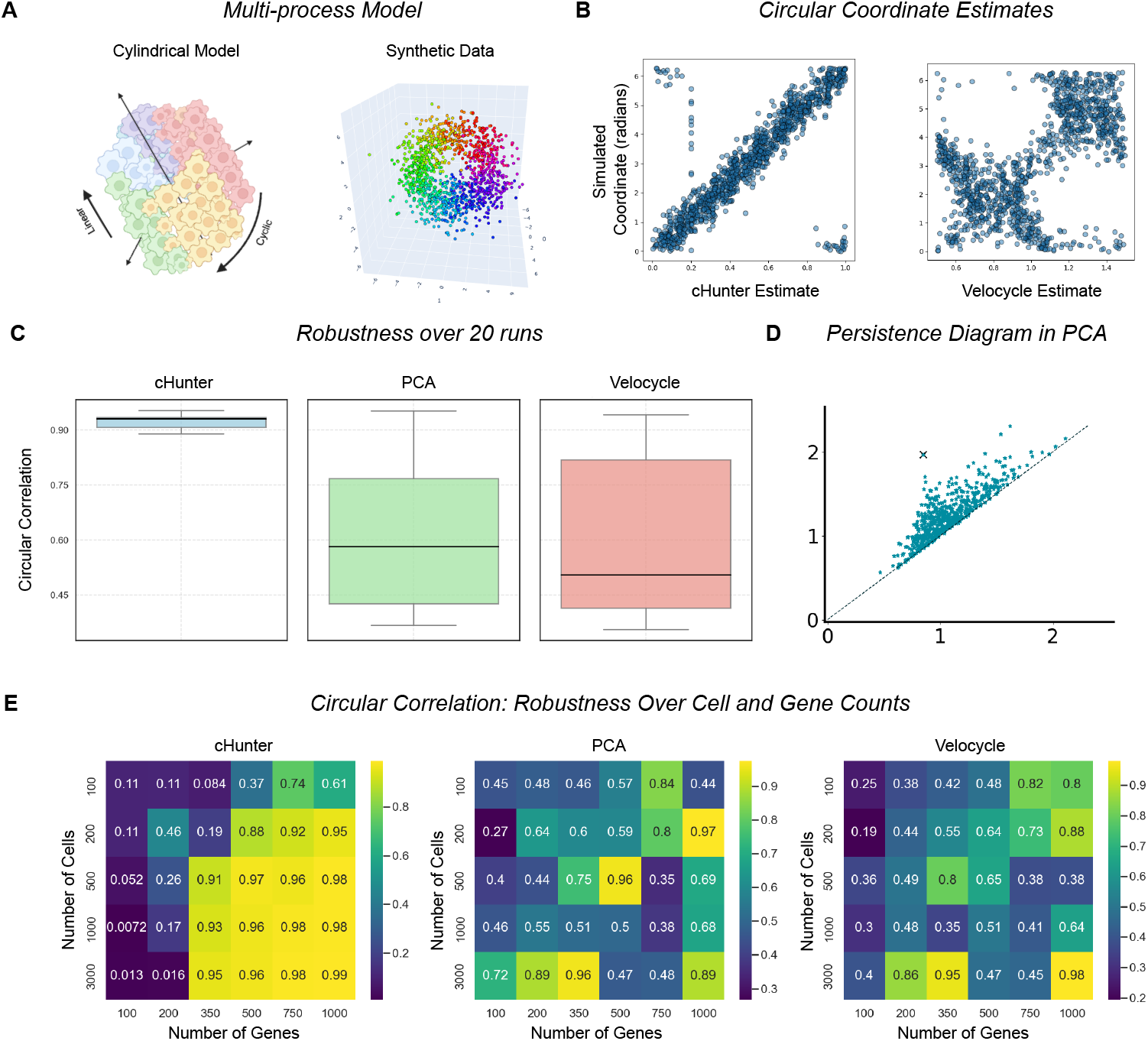
Topological methods disentangle synthetic parallel processes. **(A)** Conceptual cylindrical model and 3D PCA of the multi-process benchmark with 1200 simulated cells and 300 genes associated to each process. **(B)** Predicted circular coordinates by `cHunter` and `velocycle`. `velocycle` confuses the linear and circular processes while `cHunter` disentangles them. **(C)** Box-plot of ground-truth circular correlation of 20 instances of the multi-process synthetic data set with simulated 1200 cells, 300 circular-process genes, and 300 linear-process genes with `cHunter`, PCA baseline and Velocyle circular coordinate estimates. **(D)** Example persistence diagram of PCA 0,1,2 of such an instance. **(E)** Circular correlations with ground truth over varying cell and gene counts in the multi-process synthetic benchmark.

The topological approach taken by `cHunter` was able to reliably and robustly (Figure 3B) decouple these processes. We compared `cHunter` with a state-of-the-art method `velocycle` [8] for analyzing circular structures in single-cell data. Across various parameter settings (Figure 3E), `cHunter` consistently outperformed `velocycle` and the PCA baseline, both of which tended to conflate the circular and monotonic processes (Figure 3B). Notably, `cHunter` uses persistent cohomology to separate out circular and monotonic dynamics based on the topology of the data, whereas `velocycle` relies on a fixed single cycle prior that can conflate recurrent dynamics with other processes.

We additionally compared the estimation of circular coordinates against another, more traditional synthetic model presented in `velocycle` [8], which simulates a single cyclic process (see Appendix A.2). Sensitivity Figure S3A and robustness analysis Figure S3C over the parameters in [8] showed that `cHunter`, `velocycle` and the PCA baseline produced roughly equivalent circular coordinate predictions with the groundtruth (Figure S3A,C,D). Given that the circular structure was clearly visible in the first two principal components (Figure S3B), our analysis justified the need for more challenging settings to distinguish the approaches.

### Synthetic model shows how topology emerges from transcriptional negative feedback loops

To lay the theoretical groundwork for how topological structure can emerge from RNA regulation, we introduced a synthetic model of coordinated negative feedback loops in RNA transcription (Appendix A.1). In our model, RNA transcription was viewed as a bursty, switch-like process in which genes toggled between inactive and active states based on global gene expression, reflecting the intermittent nature of transcription observed in cells [36, 37]. Rather than assuming continuous gene expression, our approach introduced a threshold mechanism that triggered transcription only when specific conditions were met, thereby capturing the finite and episodic production of RNA. To mimic the inherent variability of single-cell data, we incorporated a sampling process that simulated the random fluctuations observed in real experiments [38–41]. When incorporating negative feedback loops into the interactions between RNA and proteins, our model naturally gave rise to cyclic patterns that were observable in low-dimensional projections (see Figure S2D). Moreover, the geometric lead-lag analysis of such cycles was able to faithfully recover the underlying ground-truth leader-follower relationships between genes S2D).

### Topology and lead-lag analysis recovers known cell cycle dynamics of human fibroblasts in culture

Firstly, we wanted to validate that our harmonic lead-lag method produced sensible results on the most commonly studied closed biological process: the cell cycle. To do so, we used a dataset of cycling human dermal fibroblasts [42] (Appendix A.3), which has been used previously as a validation dataset on cell-cycle tools [8]. We projected the data onto the gene set spanned by known S and G2M markers (we use the markers in [8]). After identifying a persistent cohomology class, the lead-lag projection (Figure 4A) exhibits coherence between the estimated phase and the transitions from S to G2M phases. Orientation was assigned based on known S and G2M orderings, and cell cycle start point was assigned minimize total counts – both choices are independent of leadlag coefficients. Further, the logarithm of the total number of molecules was consistent with passage of the cell cycle through the S phase into mitosis.

**Figure 4.**
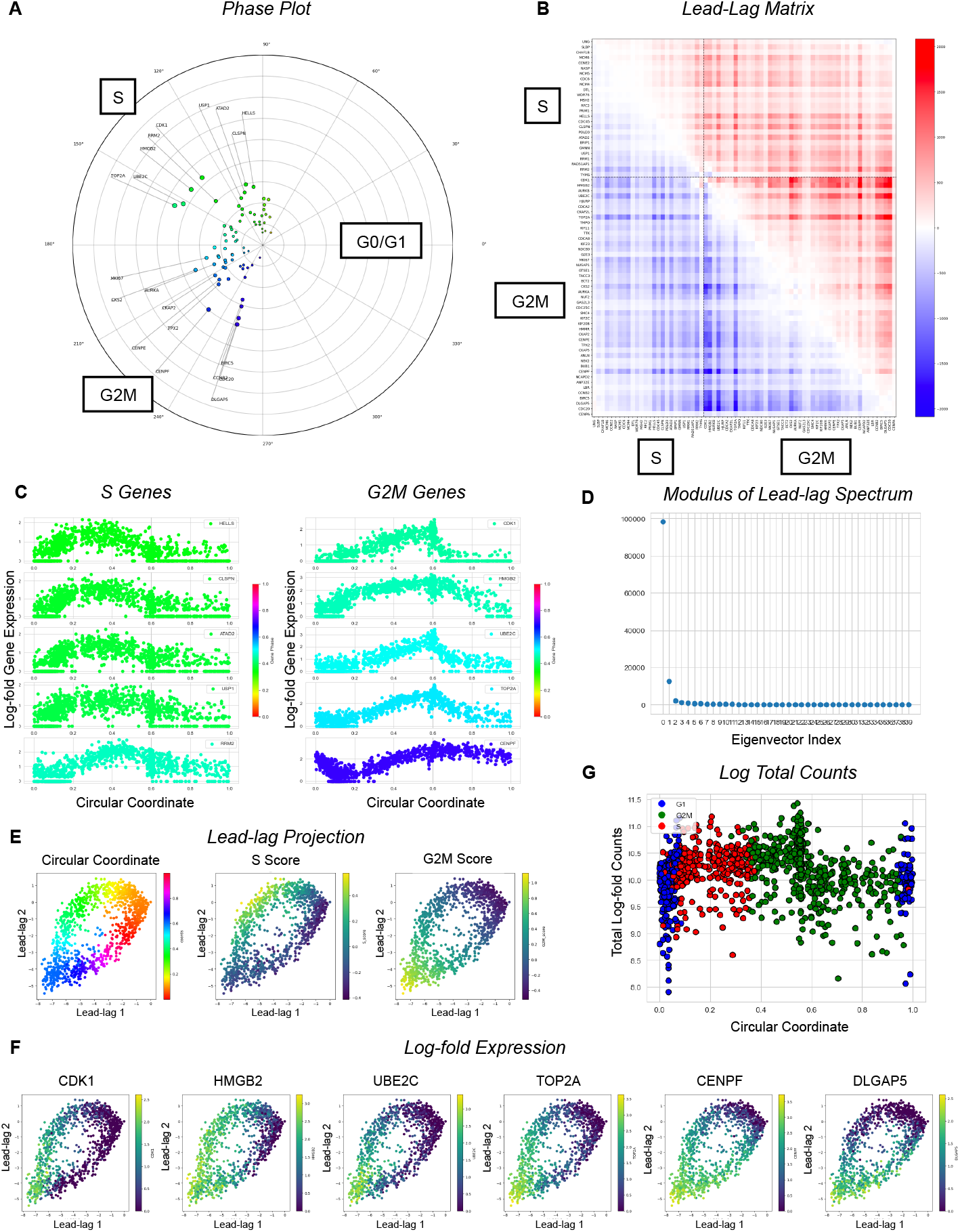
Topological methods recover known cell cycle dynamics. **(A)** Estimated gene phases from harmonic lead-lag analysis. **(B)** The lead-lag matrix, sorted by estimated gene phase and groups by *S* and *G*2*M* genes. **(C)** Normalized log-fold expression of oscillatory genes in the S and G2M phases plotted against the estimated circular coordinates. **(D)** Sorted absolute values of the complex eigenvalues of the lead-lag matrix. **(E)** The linear lead-lag projection onto the principal complex eigenplane of the lead-lag matrix colored by circular coordinates, `scanpy`-derived S- and G2M-scores. **(F)** The linear lead-lag projection onto the principal complex eigenplane of the lead-lag matrix colored by oscillatory gene expression in order of estimated phase around the cell cycle. **(G)** The log of the total reads per cell (*y*-axis) against the circular coordinates (*x*-axis).

The calculated lead-lag matrix was consistent with the high-level dynamics of S genes leading G2M genes (Figure 4D). This consistency was also present in the gene phase plots (Figure 4C) and when ordering gene expression against cell phase (Figure 4E,F). There was an observable gap of genes estimated to peak at angle 0, which again corresponds to the omission of G0/G1 genes from the cell cycle marker gene set in [8]. The human fibroblasts dataset contains the additional information of spliced and unspliced molecules. We reran our method on the same cell cycle gene set, separating out spliced and unspliced molecules. As in the previous case, we identified a salient persistent cohomology class, where the lead-lag projection was consistent with Seurat cell-cycle scoring (Figure S4A). The lead-lag matrix showed that unspliced RNA levels led spliced RNA levels of corresponding genes as expected (Figure S4A). Further, the estimated phases of unspliced molecules proceeded those of their corresponding spliced isoforms broadly across the gene set (Figure S4B,D).

### Topology shows that S and G2M cell cycle dynamics are robust under AR silencing

Existing literature suggests that the androgen receptor (AR) plays a crucial role in regulating human dermal fibroblasts (HDF) homeostasis and can influence early steps of cancer-associated fibroblast (CAF) activation. In this setting, AR can act as a key negative regulator of cellular senescence–effector genes (like CDKN1A) and of genes comprising the Senescence-Associated Secretory Phenotype (SASP) [43, 44], which includes proinflammatory cytokines and matrix remodeling proteins also found in fully established CAFs [30]. Having attained a baseline of normal cell cycle functioning in HDFs, we set out to investigate whether AR-silencing in these cells produces biologically meaningful topological and/or geometric perturbations.

Our novel single-cell RNA-seq dataset consisted of human dermal fibroblasts (HDFs) infected with two lentiviruses either expressing different AR-silencing shRNAs versus a control expressing scrambled shRNAs (Appendix A). We first aimed to compare the findings from the previous case study on fibroblast cell cycle to our second multi-conditioned (control and AR-silenced) HDF dataset. To replicate the approach, we used the same S and G2M marker genes as the previous section, all 98 of which were identified as highly variable. Lead-lag analysis on a circular structure identified in the first three principal components of this gene set revealed a striking correlation in estimated gene phases (*R* ≈ 0.96) and amplitudes (*R* ≈ 0.84) across this dataset and that of Capolupo et. al (77 shared, highly variable genes) (Fig. 5B).

**Figure 5.**
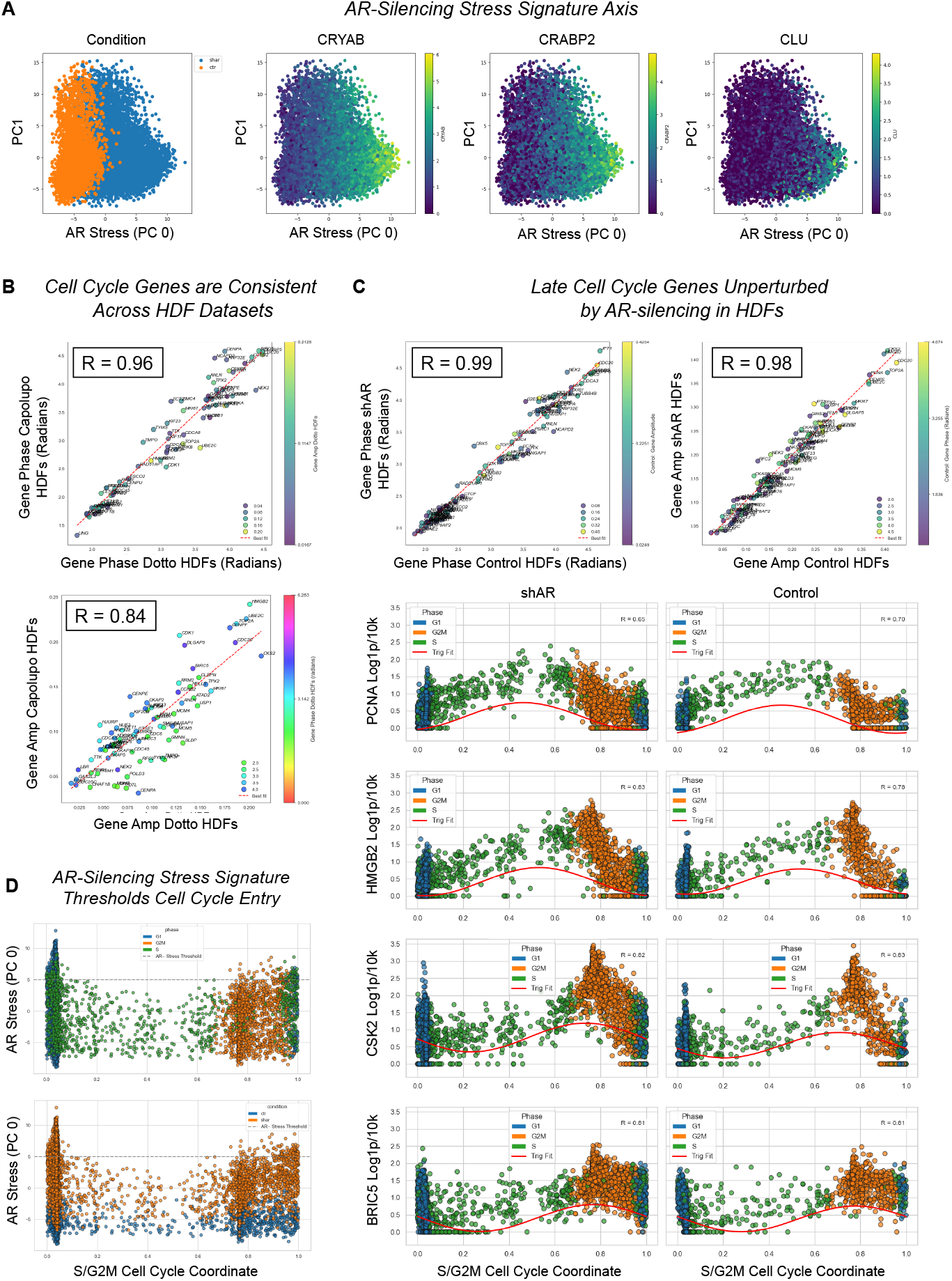
Key cell cycle genes are robust under AR silencing, but AR stress can threshold cell cycle entry. **(A)** PCA plot of first two principal components over highly-variable genes in control and AR-silenced HDFs. PC zero is taken as a continuous signature parameter indexing AR-silencing stress response. **(B)** Comparison of `cHunter` first-order gene-wise amplitude and phase estimates across HDFs in Dotto and Capolupo et al. datasets. **(C)** Comparison of `cHunter` first-order gene-wise amplitude and phase estimates across AR-silenced vs control HDFs in Dotto dataset. The expression of cell cycle genes against cell cycle coordinate in AR-silenced vs control HDFs. **(D)** AR-silencing stress response signature against cell cycle circular coordinates.

We then set about comparing the key cell cycle genes in our gene set across the two conditions within these identified cycling populations. We devised a method to re-fit normalized gene expression as a first-order sine wave over the circular coordinate using multilinear regression over the lead-lag plane. By applying this pipeline to shAR and control populations, we were able to attain separate estimates of the first-order Fourier coefficient for each cell cycle gene across conditions. We found a striking correlation between estimates of both first-order amplitude (*R* ≈ 0.98) and phase (*R* ≈ 0.99) (Fig. 5C). This suggests AR-silenced HDFs that progress through cell cycle, the core transcriptional oscillations of these key S and G2M genes are largely maintained (Fig. 5C,D). We concluded that the primary consequence of AR-silencing stress appears not to be a gross disruption of the cycle’s mechanics in still-proliferating cells, but rather an exit from the cycle in a distinct subpopulation, as explored next.

### AR stress signature decreases along S-phase of cell cycle for human dermal fibroblasts

Continuing on with the dataset consisting of AR-silenced vs control HDFs, we found that the first principal component (PC zero) captured a continuous phenotypic transition from control to shAR HDFs 5A. We termed this PC zero axis the “AR-silencing stress signature.” Crucially, when we examined this continuous signature in relation to cell cycle status, we observed that cells with high PC zero scores (indicating high AR-silencing stress) were almost exclusively found outside of the S and G2M phases, consistent with an exit from active proliferation 5D. This observation reconciles the apparently overall cell cycle distributions in bulk populations with a potent cell cycle exit occurring specifically within the high-stress subpopulation.

Examining the PC zero factor loadings provided molecular insight into this stress signature. Genes with strong positive loadings, indicative of upregulation in the high-stress/arrested state, included compelling markers of cellular senescence and the SASP. Notably, TGFBI (Transforming Growth Factor Beta Induced), a protein robustly linked to senescence and induced by TGF-*β* signaling [45], was the 5th highest-ranked contributor to this signature among the top 2000 highly variable genes, with a strong positive loading of 0.147. Further supporting this was TGFBR1, a receptor involved in TGF-*β* signaling and CAF marker [46] (rank 144, loading 0.020). The key cell cycle inhibitor and senescence marker CDKN1A (p21) [47] also exhibited a positive loading of 0.029 (rank 88). Finally, IL6, a canonical SASP cytokine that can relay and sustain senescence [44], was also highly ranked (rank 210, loading 0.014). The coordinated upregulation of these genes strongly suggests that the AR-silencing stress signature identifies HDFs undergoing G0 arrest in a state highly characteristic of cellular senescence.

### Circular enrichment identifies cholesterol homeostasis as cyclic sub-process of AR-regulated tissue regeneration ab initio

We next evaluated whether our analysis of cell cycle topology and AR function produces similar results in other settings, including (1) in vivo, (2) mouse models, and (3) across multiple scRNA-seq samples. To this end, we examined a publicly available dataset [29] where the authors study the role of androgen in prostatic epithelial cell regeneration in mice (Details A.5). In the study, mice are castrated, and within 28 days, the prostate involutes to 90% of its size. At day 28, mice are transplanted subcutaneously with dihydrotestosterone (DHT) to initiate androgen-mediated regeneration. After 28 days of regeneration, the prostate returns to homeostasis, notably characterized by quiescence and expression of the key prostate transcription factor NKX3.1. Multiple mice received the treatment concurrently, with single-cell samples of their respective prostrates taken over the full course of castration and regeneration.

Our first question was whether circular gene set enrichment was able to identify relevant biological processes. After pre-processing (See methods), we performed circular gene set enrichment using our cohomology-based ring score, testing over the 50 hallmark sets of genes [33]. We found that androgen response was the top circularly enriched subspace of statistical significance under projection onto both the first three and first two principal components (Figure 6E), validating our hypothesis that gene expression associated with androgen response would generate a closed biological loop in the singlecell phase subspace. Circular enrichment identified two other hallmark gene sets – cholesterol homeostasis and myogenesis – as ab initio gene sets with circular structure.

**Figure 6.**
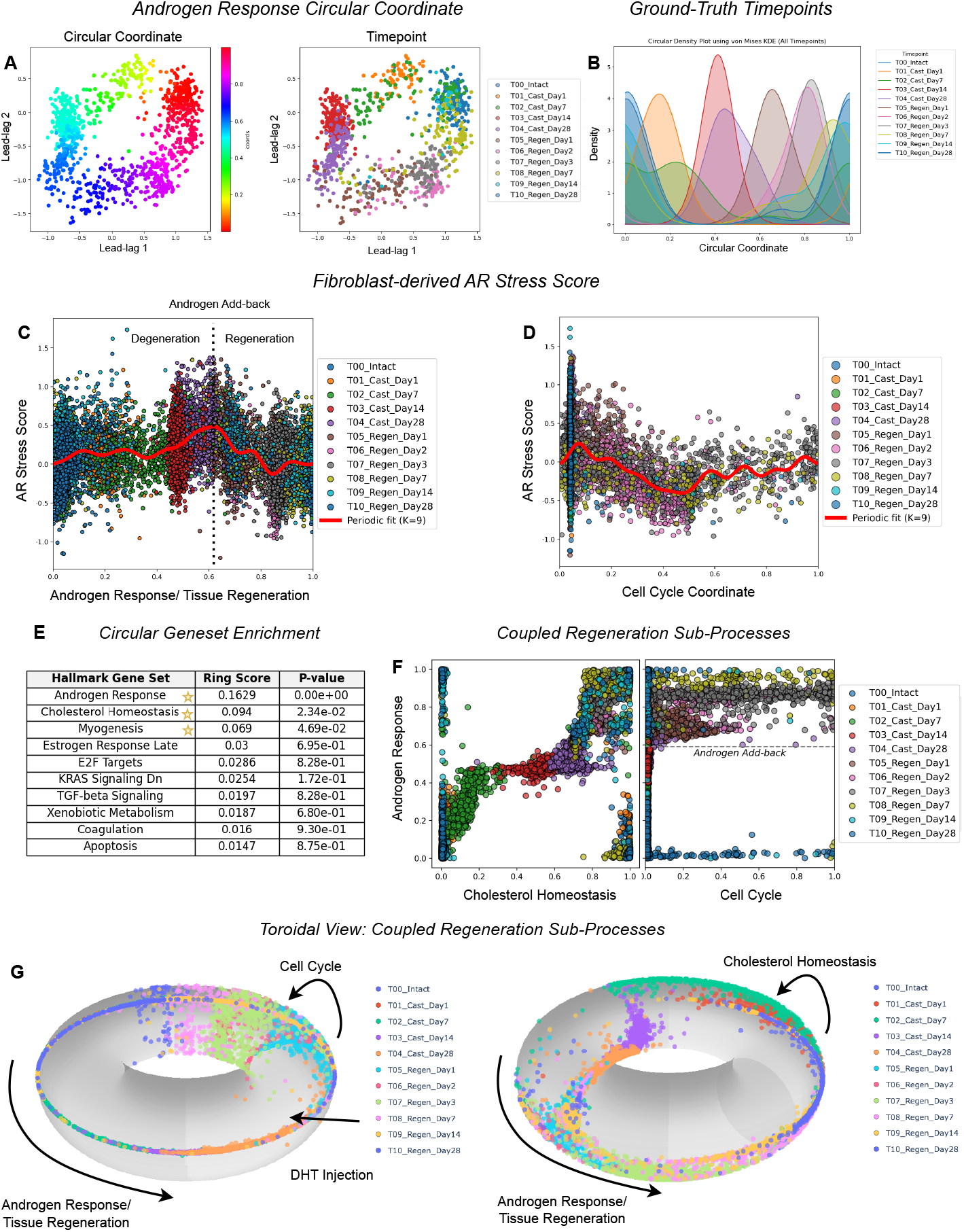
Coupling of cyclic tissue regeneration sub-processes has a topological description. **(A)** The lead-lag projection colored by androgen response circular coordinates and sample time, respectively. **(B)** The Von-Mises-based kernel density estimate of each time-point against the androgen response circular coordinate. **(C)** The androgen response coordinate against the AR stress score derived from the fibroblast data. **(D)** The cell cycle coordinate against the AR stress score derived from the fibroblast data. **(E)** The top results of circular gene set enrichment by ring score – gene sets with a p-value < 0.05 are starred. **(F)** Coupling of tissue regeneration sub-processes circular coordinates: androgen response vs cholesterol homeostasis (left) / cell cycle (right). **(G)** Toroidal view of coupling of tissue regeneration sub-processes circular coordinates: androgen response vs cell cycle (left)/cholesterol homeostasis (right).

**Figure 7.**
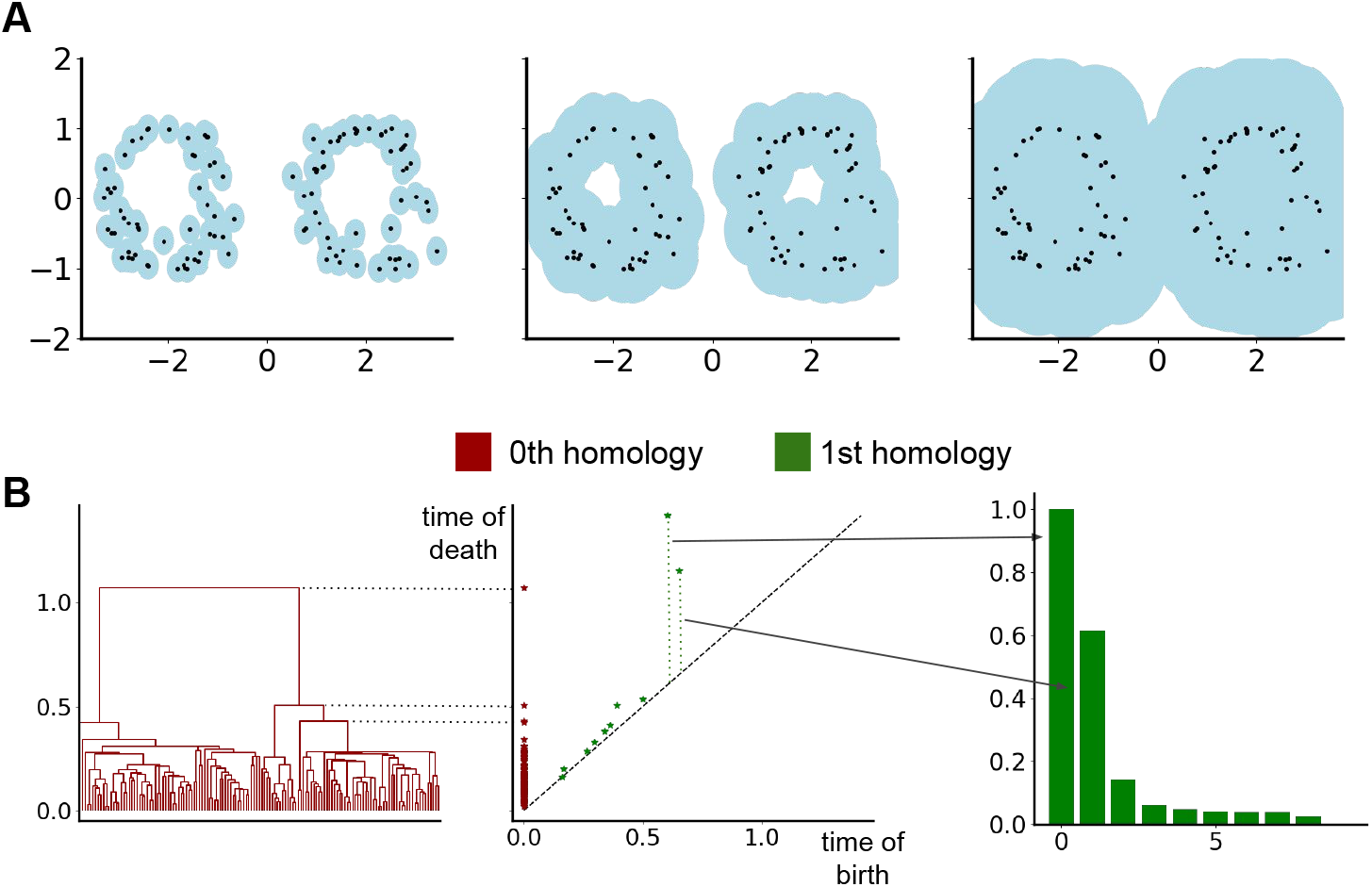
Introduction to topological data analysis. **(A)** Features that persist across a wide range of *ε* are more likely to be part of an underlying shape. Throughout the process of expanding balls around each point in a point cloud, many topological features emerge and disappear again, like connected components or “holes”. Many “short-lived” holes appear throughout this process. However, the two holes that persist the longest are more likely to be a feature of interest than the short-lived ones. **(B)** Different ways to encode persistent topological features. The middle pane is the so-called *persistence diagram*, in which each dot represents a topological feature described by its time of birth and time of death. For 0-dimensional homology classes a dendrogram is a common data structure to keep track of merging clusters (left pane). For 1-dimensional cohomology classes we consider the *persistence sequence*, i.e, the sequence of differences between birth and death times of the persistent classes, ordered by length. The first two bars indicate that there are two clear “signals” of cycles, whereas the rest of the bars indicate cycles that are considered to be “topological noise”.

The experiment provided a ground-truth labeling of samples ordered by time in the underlying regeneration process. These labels were respectively ordered by: intact; 1 day, 7 days, 14 days after castration; 1 day, 2 days, 7 days, 14 days and 28 days after regeneration. Without using explicit knowledge the timepoints in the computation, the estimated circular coordinate associated to androgen response gene set was coherent with the ground truth ordering of the samples (Figure 6B). We concluded that the circular structure found in the androgen-response gene set was an effective continuous parametrization of the tissue regeneration process, and highlights that temporal processes can be estimated directly from the geometry of the data.

We then performed the associated lead-lag analysis (Figure S6A-C). We noted that several androgen-regulated genes in the matrix have key roles in prostate development and maintenance. Notably, NKX3.1, a transcription factor known to be essential for prostate development and maintenance of luminal cell [48] identity, was estimated to peak at homeostasis time-points before castration and after full regeneration. We also noted the high amplitude of the androgen-regulated SGK1, which plays a role in cellular stress response. Lastly, we observed a strong effect of androgen-regulated cell-cycle gene gets in the early phase of regeneration, in line with the high proliferation observed. Of note, these androgen-response genes have been generated in prostate cancer cell lines that are dependent on androgens for growth.

The experiment provided a ground-truth labeling of samples ordered by time in the underlying regeneration process. These labels were respectively ordered by: intact; 1 day, 7 days, 14 days after castration; 1 day, 2 days, 7 days, 14 days and 28 days after regeneration. We observed that the estimated androgen response circular coordinate was coherent with the ground truth ordering of the samples (Figure 6B), despite being calculated independently of this information. We concluded that the circular structure found in the androgen-response gene set was an effective continuous parametrization of the tissue regeneration process.

In addition to the androgen-response gene set, circular gene set enrichment identified the cholesterol homeostasis gene set as statistically significant with respect to the ring score (Figure 6E). Plotting the circular coordinates against each other (Figure 6F,G) showed that both circular coordinates were dependent on each other, which highlights that these biological sub-processes associated with the master tissue regeneration cycle are strongly coupled. Potentially, this is an intrinsic adaptation to the absence of androgens, as cholesterol is a precursor to androgens. Additionally, the union of the gene sets also exhibited a dominant circular structure, from which the lead-lag analysis allowed the ordering of the genes between the sets to be compared. This demonstrated that the two processes are dominant at different points of the process, as per the phase plot in Figure S6G.

### Human fibroblast-derived AR-silencing stress signature functions consistently in regenerating mouse epithelia

Finally, we sought to compare whether the transcriptional AR-deprivation response was governed by comparable genes across HDFs and prostatic mouse epithelial cells. We observed that our HDF-derived AR stress signature was maximally valued on prostatic epithelial cells at the peak of androgen deprivation, 28 days after castration, before gradually decreasing after androgen add-back (Figure 6C and Figure 6K). This suggests that our AR stress signature was measuring a consistent transcriptional response related to AR in two settings: (1) where HDFs are AR-silenced in culture and (2) where mouse prostate epithelial cells are deprived of androgen via castration.

We also wanted to examine the relationship between AR stress signature in the prostate data. Interestingly, cell cycle gene sets were not found to be circularly enriched in the prostate data. We were, however, able to identify a persistent cohomology class corresponding to the cell cycle by restricting to the first 7 days of regeneration and the 28th day after castration, suggesting that the geometry of the tissue regeneration cycle obscures the cell cycle in the full dataset. Analogous to the HDFs, androgen add-back mediated progression through the S-phase of the cell cycle Figure 6F,G was associated with a lower AR stress signature, suggesting a potential common transcription pathway across human and mouse cell-types regulating cell-cycle progression and AR-signaling.

## Discussion

Our work introduces a principled topological-geometric framework that detects and disentangles multiple concurrent cyclic processes in single-cell transcriptomic data. By relaxing the traditional one-dimensional manifold assumptions that dominate current single-cell analysis pipelines, we demonstrate that topological and geometric techniques are capable of capturing the multidimensional and recurrent structure of biological dynamics. This approach provides a unified mathematical formalism to recover and interpret cyclic processes that are typically conflated or obscured by standard clustering or trajectory inference methods.

The identification of multiple, concurrent cycles as separate circular dimensions in single-cell data highlights a fundamental limitation of Waddington’s classical landscape metaphor. While the landscape model elegantly describes differentiation as a branching and irreversible process, it lacks the formalism to represent concurrent cyclic phenomena that govern many biological systems, such as the cell cycle and tissue regeneration. Our results establish that the transcriptomic manifold of single cells is not simply tree-like, or even circular, but is better described by a richer collection of higher-order topological dimensions—circles, tori, and potentially more complex substructures—that reflect the coexistence of multiple biological clocks. The example in Figure 6G demonstrates a key observation about the Waddington landscape: it is *multi-temporal*, i.e., it exhibits multiple dimensions corresponding to temporal processes.

We first validated our method using synthetic datasets where we designed a cylindrical model of concurrent linear and cyclic processes. In this controlled setting, our framework—implemented as the `cHunter` software package—robustly separated circular from linear dynamics, outperforming state-of-the-art methods that assume a single global cycle. This benchmark confirmed that topological methods generalize beyond prior heuristics by allowing the data itself to define the number and structure of recurrent processes. Importantly, our synthetic modeling of negative transcriptional feedback loops provided mechanistic intuition for how cyclic topology can emerge naturally from gene-regulatory interactions, linking geometric structure to biological causality for scRNA-seq data.

Focusing on primary human dermal fibroblasts, we validated the use of topology and lead-lag analysis to detect circular cell and gene transcription dynamics across the cell cycle, with consistent estimates underscoring the robustness of our approach. The harmonic coordinates and lead–lag matrices recovered coherent transitions from S to G2/M phases, faithfully reproducing known cell-cycle gene relationships. We found that cells responded heterogeneously to silencing of the AR gene, and that AR-stress–related cell-cycle arrest coincided with increased expression of key senescence-related genes, including TGFBI, CDKN1A, and IL6. The relationship between our AR-stress signature and cell-cycle obstruction was observed in multiple settings—in vivo mouse epithelia and in vitro human fibroblasts—suggesting a shared underlying molecular mechanism. These findings are consistent with a continuous phenotypic transformation from normal stromal fibroblasts to cancer-associated fibroblasts (CAFs), a process that may be reversible through epigenetic reprogramming.

Our topological approach of circular gene set enrichment revealed cholesterol homeostasis as a key process in AR-regulated tissue regeneration. This previously unrecognized connection has significant biological implications, including the involvement of enzymes shared between cholesterol and androgen biosynthesis, which could be therapeutically targeted in disease contexts such as prostate cancer or stromal reprogramming. This example demonstrates the viability of circular enrichment as a generic topological technique for the unbiased, de novo identification of gene sets associated with cyclic transcriptional processes. In the in vivo prostate regeneration model, the harmonic ordering of androgen-response and cholesterol-homeostasis gene sets suggested that these processes are interdependent sub-processes of a broader regenerative program, supporting the concept of nested cyclic hierarchies within tissue-level regeneration.

Through these analyses, we demonstrated the applicability of our ab initio topological approach across diverse settings—including synthetic RNA models and benchmarks, in vitro human cells, and heterogeneous in vivo mouse data. We have introduced topological and geometric statistics to estimate the multidimensional geometry of the Waddington landscape directly from data, extending beyond clustering and single-parameter methods. This framework allows practitioners to isolate multiple cyclic processes and study their interactions, capturing both irreversible and recurrent dimensions of cellular state transitions. We envision that this represents a first step toward a general geometric theory for factorizing dynamic transcriptomic phenomena into simpler subprocesses—or “process times” [49]—and for estimating the underlying multi-temporal topological structure of the Waddington landscape.

## Funding acknowledgements

K.M., M.K.Y and J.I. were supported by the European Union’s Horizon 2020 Research and Innovation Program under Marie Skłodowska-Curie Grant Agreement No 859860. W.K was supported by Swiss Cancer League grant number KLS-5654-08-2022 and Swiss National Science Foundation, project number: 310030-215517. Research in the G.P.D laboratory was supported by grants from the Swiss National Science Foundation (310030B_176404 “Genomic instability and evolution in cancer stromal cells”) and the NIH (R01CA269356, the contents do not necessarily represent he official views of the NIH). K.M., M.A. and H.A.H. are members of the Max Planck/Oxford Center to Center collaboration supported by the Mathematical Foundations of Intelligence: An “Erlangen Programme” for AI, reference: EP/Y028872/1. K.M. and H.A.H. are members of the EPSRC Erlangen programme for AI Hub EP/Z531224/1.

## Methods

### Single Cell RNA-seq

#### Basic Background

Single-cell sequencing is a technology that measures the number of RNA molecules in each cell in a given tissue. For our purposes, the mathematical output of a single-cell experiment consists of a matrix `expression` whose rows correspond to the cells and columns to genes. Equivalently, the matrix corresponds to an integer point cloud whose points are cells and dimensions are genes.

#### Pre-processing

Single-cell data is well-known to be noisy. There are several common pre-processing steps to filtering and processing the initial expression matrix that we apply in our experiments. Following the Seurat [1] recommendations, our pipeline is as follows:

1. Filter genes and cells by a minimal number of molecules.
2. Library normalization and variance stabilization. In our case, we used an ℓ_1_-normalization scaled by 10000, followed by a log-transformation, as suggested in [2]

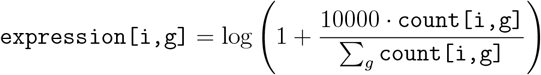
3. We optionally apply a diffusion scheme using the MAGIC^1^ algorithm [3].
4. Taking subsets of the remaining gene set determined by highly variable genes.

### Mathematical background

#### Persistent cohomology

##### Simplicial complexes

A (finite) *simplicial complex* is a (finite) set *X* and collection 𝒮 of subsets of *X* satisfying

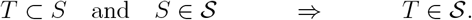

The sets *S* ∈ 𝒮 are *simplices* and sets *S* ∈ 𝒮^*k*^ of cardinality *k* +1 are *k-simplices*. If *X* has *n* elements, the *geometric realization* of (*X*, 𝒮) is topological subspace

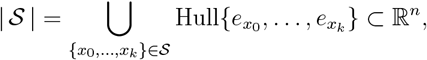

where to each element *x* of *X*, we associate a different choice *e*_*x*_ of standard basis vector of ℝ^*n*^, and Hull denotes the convex hull. A simplicial complex is *oriented* if there is a total order on *X*, in which case simplices are written as ordered tuples

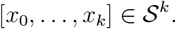

##### Cochains

Let (*X*, 𝒮) be an oriented simplicial complex and *R* a commutative ring. The *k-cochains* of (*X*, 𝒮) is the *R*-module

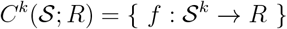

of *R*-valued functions on the set 𝒮^*k*^ of *k*-simplices. The *k-th coboundary operator* is *R*-linear map

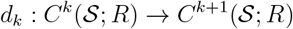

defined by

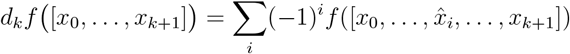

where 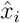 is omitted from [*x*_0_,…, *x*_*k*+1_]. The *simplicial cochain complex C*^•^(𝒮; *R*) of (*X*, 𝒮) is the sequence of *R*-modules

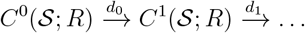

and satisfies the relation *d*_*k*+1_*d*_*k*_ = 0 for all *k*. The *k-th cohomology* of the complex is given by the quotient

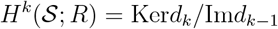

of the *k-cocycles* Ker*d*_*k*_ modulo the *k-coboundaries* Im*d*_*k*−1_. A map *φ*: (*X*, 𝒮) → (*Y*, 𝒯) is *simplicial* if *φ* (*S*) ∈ 𝒯 for all *S* ∈ 𝒮. If *φ* is simplicial, then it induces an *R*-module homorphism on cohomology

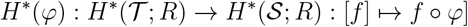

for any ring *R*.

##### Persistent cohomology

Let (*X, d*) be a metric space, i.e., a set *X* equipped with a metric (distance) *d*. The *Vietoris-Rips complex* at distance *ε* ∈ ℝ_+_ is the simplicial complex *X*_ε_ with *k*-simplices

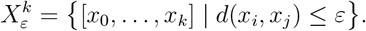

For each *ε < δ*, there is an inclusion *X*_ε_ ↪ *X*_*δ*_, thus given 0 = *ε*_0_ *< ε*_1_ *<*…*< ε*_*n*_, we obtain the *Rips filtration*

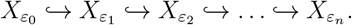

The *k-th persistent cohomology H*^*k*^(*X*_*•*_) of (*X, d*) over a field 𝔽 is then the sequence

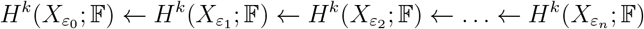

obtained from applying cohomology to the Rips filtration.

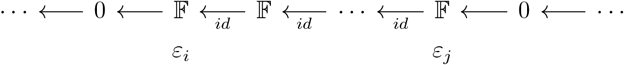

The following theorem forms the basis of practical applications of persistent cohomology, and is attributed to different references depending on the conditions [4–6]. It is formulated in terms of *intervals* 𝕀_[ε,*δ*]_, for *ε* ≤ *δ*, which is the following sequence of 𝔽-vector spaces and linear maps.

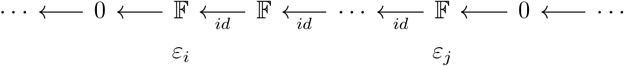

###### Theorem 1

(Interval decomposition). *For each k, the persistent cohomology H*^*k*^(*X*_*•*_; 𝔽) *of a finite metric space* (*X, d*) *is isomorphic to a direct sum of interval modules*

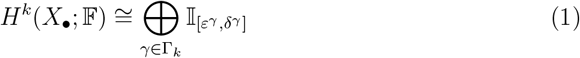

*which is unique up to permutation of summands*.

The intervals 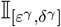 in the decomposition are called *persistent cohomology classes*. The start *ε*^γ^ and end *δ*^γ^ points are the *birth* and *death* times of the persistent cohomology class. The *persistence diagram* of (*X, d*) is the multi-set

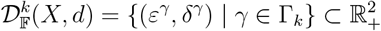

using the interval decomposition in Equation (1).

##### Stability

The *bottleneck distance* between two persistence diagrams is

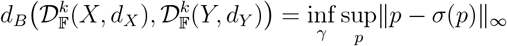

Where 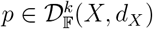, and *σ* ranges over all bijections 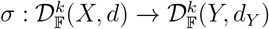. Let

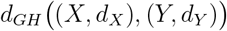

denote the Gromov-Hausdorff distance between two finite metric spaces (*X, d*_*X*_) and (*Y, d*_*Y*_) (see [7] for the definition). The following theorem states that the persistence diagram is stable with respect to small perturbations of the underlying metric space.

###### Theorem 2

([7], Thm. 3.1).

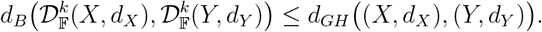

#### Harmonic circular coordinates

In this section we review the circular coordinates algorithm from [8], emphasizing the density-weighted [9] and sparse [10] variants.

##### Representability

For any pair *X, Y* of pointed topological spaces, let [*X, Y*] _∗_denote the set of pointed homotopy classes of basepoint-preserving continuous maps from *X* to *Y*. The following classical theorem provides an important interpretation of cohomology over ℤ. Let *S*^1^ denote the usual circle of radius 1 in the plane, centered at the origin.

###### Theorem 3.

*(cf. Thm. 4*.*57 [11]) For any simplicial complex* 𝒮, *there is a bijective correspondence*

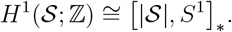

##### Harmonics

The bijective correspondence above between cohomology classes in degree 1 and homotopy classes of maps into the circle begs the question of which map to pick within the homotopy class to best represent the cohomology class. A particularly useful approach begins with the choice of a family of inner products

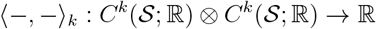

on the cochain complex. The *combinatorial Laplacian* is then the linear map

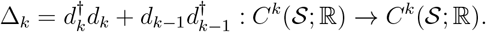

where 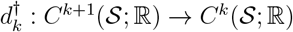 is the adjoint map satisfying

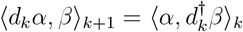

for all *α, β*. According to he fundamental theorem of Hodge theory, proven for the simplicial case in [12], there is an isomorphism

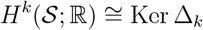

between the *space of harmonics* KerΔ_*k*_and the cohomology. The space of harmonics depends on the choice of inner product and consists of the *k*-cochains that minimize the induced norm within their cohomology class.

##### Simplicial circular coordinates

A simplicial cochain *α* ∈ *C*^1^(𝒮; ℝ) has *integral periods* if there exists a integral cocycle *α* _ℤ_∈ *C*^1^(𝒮; ℤ) such that

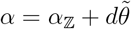

for some 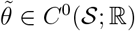.In this case, there is a well-defined function

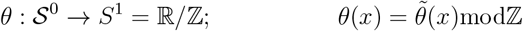

Satisfying

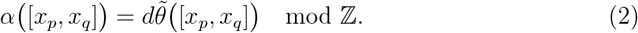

The map *θ* extends to a continuous map defined on the geometric realization |𝒮| → *S*^1^ (see [8]).

##### Harmonic smoothing

Equation (2) implies that the harmonic representative

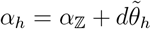

corresponds to a circular coordinate *θ*_*h*_: 𝒮^0^ → *S*^1^ with the ‘smoothest’ gradient (depending on choice of inner product). This is attained as a solution to the Dirichlet equation

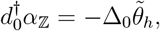

which is computed via least-squares regression. We refer to the associate continuous map *θ*_*h*_: |𝒮| → *S*^1^ as the *circular coordinate associated to α*_ℤ_and refer the reader to [8, 9] for further details.

###### Remark 4

*The map θ*_*h*_*depends only on the inner products in dimensions* 0 *and* 1.

##### Persistence-based coordinates

Persistent cohomology algorithms are computed over cofficients in the field ℤ*/p* for some prime *p*. This issue is mitigated by the following observation from [8].

###### Proposition 5

*For a finite simplicial complex* 𝒮, *the map*

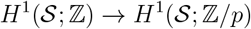

*induced by the unique ring map* ℤ → ℤ*/p is surjective whenever H*^2^(𝒮; ℤ), *or equivalently H*_1_(𝒮), *has no p-torsion*.

Since torsion is remarkably rare in data, we can assume that we have a valid lift of a (ℤ*/p*)-class to a ℤ-class. For an interval 𝕀 _[ε,*δ*]_in *H*^1^(*X*_*•*_; ℤ_*p*_), we lift the cocycle representative *α* ∈ *C*^1^(*X*_ε_; ℤ_*p*_) to a ℤ-valued cocycle *α*_ℤ_∈ *C*^1^(*X*_ε_; ℤ). The corresponding circular coordinate is the harmonic circular coordinate

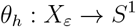

corresponding to the harmonic representative 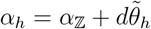. See [10] for details.

##### Density weighting

The single-cell data we work with has highly non-uniform density in practice. In [9], the authors define cochain inner products that correct for density and metric information, following the graph-Laplacian manifold-approximation literature [13].

For a finite point cloud (*X, d*) in ℝ^*n*^, define the *kernel* and *density* functions as

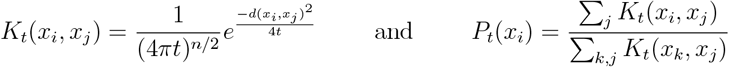

for a parameter *t* ∈ ℝ_+_. For a cut-off distance *ε* ∈ ℝ_+_, let

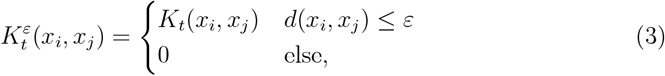

and define 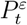 similarly. Define the *kernel-density weighting W*: *X* × *X* → ℝ by

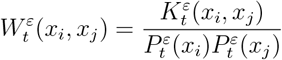

The inner products on *C*^0^(𝒮; ℝ) and *C*^1^(𝒮; ℝ) given in [9] are defined via the formulae

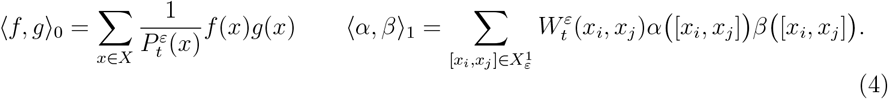

###### Remark 6

*The inner products correspond to density-normalized correlations of* 0*and* 1*-cochains*.

When *X* is sampled from a distribution on an embedded Riemannian manifold, the combinatorial Laplacian Δ_0_under these inner products approximates (for an appropriate choice of *t*) the intrinsic Laplace-Beltrami operator, irrespective of distribution (Thm. 3.1 [9]).

##### Sparse circular coordinates

In practice, one mitigates the computational complexity of persistent cohomology by working over landmark subsets [10]. Briefly, a landmark set

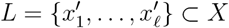

is attained by inductive maxmin sampling

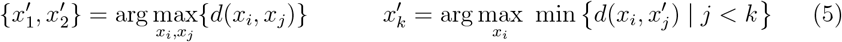

until reaching a specified distance

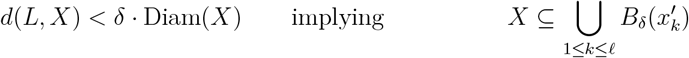

for a parameter *δ* ∈ ℝ_+_(we use 0.01 by default). Assuming the longest 1-interval 𝕀_[ε,ε_]*′* in *H*^1^(*L*_*•*_; ℤ_*p*_) satisfies *δ < ε*, we compute the harmonic circular coordinate

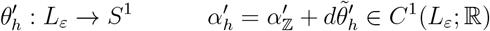

for *L* at resolution *ε* using the inner products from Equation (4). Following [10], the circular coordinate is extended from *L* to *X* by the formula

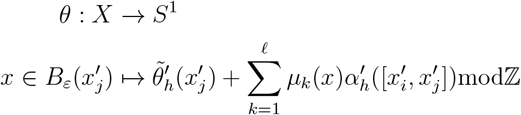

where 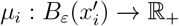 is a partition of unity. This is well-defined, independent of the choice of covering neighbourhood for each *x* (see [10] Thms 3.2 and 4.2).

#### Circular Gene Set Enrichment

Gene set enrichment analysis (GSEA) [14] is a mainstay of single-cell RNA-seq workflows. We extend this idea to *circular enrichment*: a gene set is *circularly enriched* in a cell population when the expression profile restricted to that set forms a statistically significant circular structure.

##### Gene set projections

Let *X* denote a single-cell RNA-seq data matrix that has already been fully pre-preprocessed (normalization, noise removal, and restriction to a single cluster or cell type).

###### Definition 7

*For a gene set* 𝒢 = *{g*_1_,..., *g*_*ℓ*_*} define:*

1. *X*_𝒢_: *the restriction of X to the coordinates indexed by* 𝒢;
2. 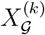: *the projection of X*_𝒢_*onto its first k principal components*.

Note that the matrices *X, X*_𝒢_, and 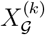 contain the same cells; only the ambient geometry changes. Intuitively, *X*_𝒢_isolatethe biological program associated with 𝒢, while 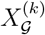 offers a lower dimensional representation of that program. Throughout this work, we set *k* = 3 — large enough to capture the dominant variation yet small enough to suppress noise. By examining 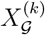, we assess whether the dominant structure of this gene set manifests as a circular pattern.

##### Ring score formula

To efficiently screen large collections of gene sets, we introduce the *ring score* — a single statistic that quantifies how strongly the subspace 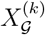 exhibits circular topology. Such ring-like geometry is characteristic of cyclic programs such as the cell cycle or circadian rhythm.

A point cloud is called *ring-structured* when it contains one prominent one-dimensional hole, as in (a sampled) circle, annulus, or hollow cylinder. The ring score, therefore, gauges how closely a point cloud resembles these ideal shapes. We derive it from the one-dimensional persistence diagram of 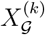.

Let the persistence diagram consist of *n* birth-death pairs

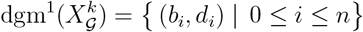

and denote their lifetimes *p*_*i*_= *d*_*i*_— *b*_*i*_, where the indices are ordered such *p*_0_≥ *p*_1_≥.…The dominant lifetime *p*_0_reflects the main circular feature, whereas the remaining lifetimes represent additional loops that “blur” this signal.

###### Definition 8

(Generalized Ring Score Formula) *Let* **p** = (*p*_*i*_)_*i*∈**N**_*be a non-increasing sequence of non-negative real numbers and choose a normalization constant N* ≥ *p*_0_. *Then we define the* generalized ring score formula *as*

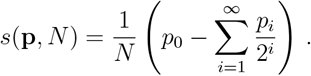

###### Remark 9

*Because* 0 ≤ *p*_*i*_≤ *p*_0_*for all i and* 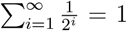, *the range of the score is given by s* ∈ [0, 1], *with larger values indicating a more dominant circular structure. The dyadic wights* 2^—*i*^ *dampen the influence of less persistent features and, together with the normalization by N, ensure that the score lies within the unit interval. More formally*,

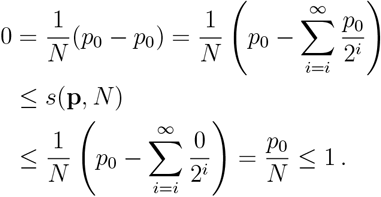

##### Scale invariance of ring score

Let 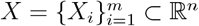 be a finite point cloud with the center of mass 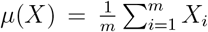 *Metric scaling* by a factor *c* > 0 keeps the centroid fixed while multiplying every pairwise distance by *c*:

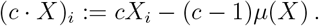

Write **p**(*X*) = (*p*_*i*_)_*i*∈ℕ_for the ordered lifetimes in the one-dimensional persistence diagram of *X*, and let *N* (*X*) be the normalization constant that appears in Definition 8. If *N* (*X*) is chosen so that

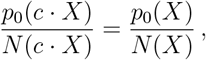

then the ring score is invariant under metric scaling. In particular, two natural and convenient choices for *N* satisfying this requirement are:

- *N* (*X*) = *p*_0_(*X*), the longest lifetime itself;
- *N* (*X*) = diam(*X*), the Euclidean diameter of the point cloud.

In fact, for point clouds embedded in ℝ^*n*^ the ratio 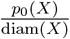 achieves its theoretical maximum of 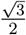, when *X* is isometric to a perfect circle.

##### Two families of ring scores

The ring score formula of Definition 8 merely requires a non-increasing sequence of non-negative numbers and in particular this sequence does not need to be persistence lifetimes. An increasingly popular alternative to lifetimes is to use *persistence ratios*, 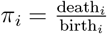.

Because no shape maximizes *X*), a convenient scale-invariant choice is simply *N* (*X*) = *π*_0_(*X*). Thus we define the following two families of ring scores.

###### Definition 10

(Diameter and ratio ring scores) *Let X* ⊂ ℝ^*n*^ *be a finite point cloud with diameter d* = diam(*X*). *Denote by* **p** *and by* ***π*** *the sequences of lifetimes and persistence ratios, respectively. We define*

- *the* diameter ring score *as* 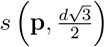.
- *the* ratio ring score *as s*(***π***, *π*_0_).

##### Iterative Density Filtering

While persistent homology is robust under perturbations to the underlying point cloud, it can be somewhat sensitive to outliers. To mitigate this, we elected to perform a basic kernel-density filtering as a pre-processing step within the circular enrichment pipeline. For a choice of bandwidth parameter *σ* ∈ ℝ_+_, we used the Euclidean kernel

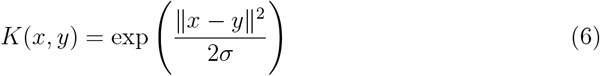

over points *x, y* in Euclidean space ℝ^*n*^. We used the quantity

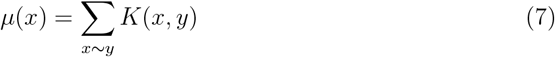

where *x* ~ *y* if *y* is one as a rough estimate of the density function on the point. We then subset the data using upper and lower percentiles of the density function in Equation (7), repeating this process for a fixed number of times *n*.

##### Ring score significance estimation

We estimate the statistical significance of ring scores in terms of nominal *p*-value, using an empirical gene set-based permutation test procedure that preserves the correlation structure of the gene expression data. For any observed gene set 𝒢^obs^ of size *ℓ*, we randomly generate a gene set 𝒢 of size *ℓ* to create and recompute ring scores of the random gene sets, generating a null distribution *S*_NULL_for the ring score. The empirical *p*-value of the observed ring score is then calculated relative to this null distribution. We note that the permutation of gene labels preserves cell-cell correlations and, thus, provides a more biologically reasonable assessment of significance than would be obtained by permuting both genes and cells.

Explicitly, given an observed gene set with *ℓ* genes 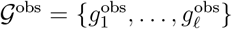 and preprocessed data *X*, we calculate significance of the ring score 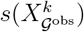 compared to a set of scores *S*_NULL_(*k, ℓ*) computed with randomly selected gene sets of size *l* in three steps.

1. Randomly select *l* distinct genes, 𝒢 = *{g*_1_,..., *gℓ}* and compute 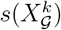.
2. Repeat step 1 for 2^10^ randomly generated gene sets, and create a null-distribution of the corresponding ring scores *S*_NULL_(*k, ℓ*).
3. Use *S* (*k, ℓ*) to estimate the nominal *p*-value for 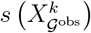.

This procedure is conceptually similar to estimating *p*-values for gene set enrichment analysis. [14]

### Cyclicity Analysis

#### Path cyclicity analysis

Cyclicity analysis is a method introduced by [16] to analyse the interaction between periodic paths in ℝ^*n*^ and their embedding dimensions. In this section, we provide an overview of the definitions and basic theory.

##### Lead-lag matrix

Let *γ*: *S*^1^ → ℝ^*n*^ be a closed path. The *lead-lag matrix* of *γ* is the *n* × *n* matrix

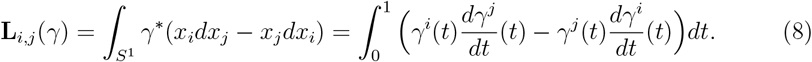

The (*i, j*)-entry of this matrix is called the (*i, j*) *lead-lag coefficient*. In simple models (cf. (12)), this coefficient measures the extent to which coordinate *i* leads or lags behind coordinate *j*. Following from the properties of integration, the lead-lag matrix is

1. skew-symmetric, and
2. invariant under orientation-preserving reparametrization of *γ*.

If 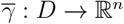 is differentiable map of a disk into ℝ^*n*^ such that the composite

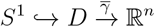

is equal to *γ*, then Stoke’s theorem implies that

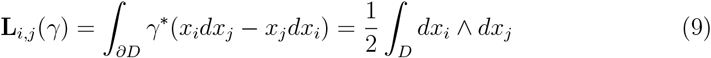

i.e., the lead-lag is proportional to the signed area of the disk in the (*ij*)-plane in ℝ^*n*^. It follows that the lead-lag matrix is invariant under translation of *γ*.

##### Chain-of-offsets model

For intepretability, [16] presents the following model. Suppose χ: *S*^1^ → ℝ is a generating periodic function with Fourier decomposition given by the real part of

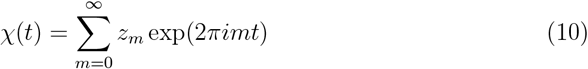

for complex Fourier coefficients *z*_*m*_∈ ℂ. A path *γ*: *S*^1^ → ℝ^*n*^ is a *chain-of-offsets model* if there exist **a** ∈ ℝ^*n*^ and **b** ∈ [0, 1)^*n*^ such that the components satisfy

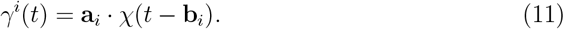

In terms of biology, the chain-of-offsets model assumes that gene expression values in different coordinates around a circular expression pattern are offset copies of one another — up to a reparametrization.

##### Lead-lag of the chain-of-offsets

The upside of the chain-of-offsets model is that the lead-lag matrix has the following closed form, due to [16]. Suppose that *γ*: *S*^1^ → ℝ^*n*^ has offsets as in Equation (11) and that the generating function χ: *S*^1^ → ℝ has Fourier decomposition in Equation (10).

###### Proposition 11

(Prop. 5.1., [16]) *The lead-lag matrix of γ is*

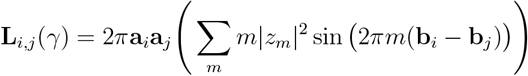

##### Ellipse model

Following [16], suppose we have a simple case where the primary function

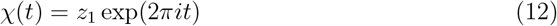

has a Fourier series concentrated in the first harmonic, i.e., that the path is an ellipse centered about the origin. In this case, applying Proposition 11 to the chain-of-offsets *γ*: *S*^1^ → ℝ^*n*^ shows that the lead-lag matrix is

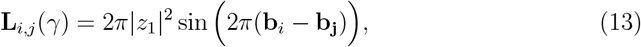

we can derive an important interpretation:

*the* (*ij*)*-leadlag coefficient is proportional to the sine of the difference in offsets of dimension i and j*.

Such a quantity is maximized and minimized when coordinate *i* is offset by a quarter and a negative quarter of the total period from coordinate *j*, respectively. As is argued in [16], if the Fourier series of χ (*t*) is dominated by *z*_1_, i.e., if

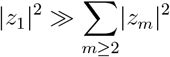

then

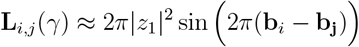

provides a similar interpretation of the lead-lag coefficients up to noise.

##### Complex diagonalization

Motivated by the fact that the lead-lag matrix is skew-symmetric, we review some basic results about the spectral theory of skew-symmetric matrices.

Let **A** be a real *n* × *n* skew-symmetric matrix. A basic result from linear algebra is that there exists a real orthonormal matrix **P** ∈ *O*(*n*) and a block diagonal *n* × *n* matrix

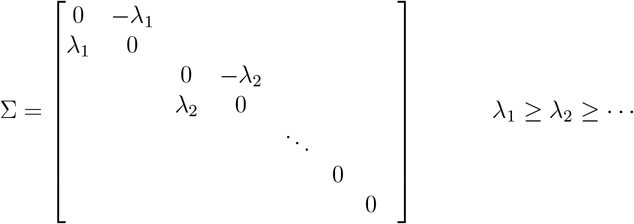

satisfying **A** = **P**^—1^∑**P**. Equivalently, treating **A** as an *n* × *n* complex matrix, there exists a unitary matrix **U** ∈ *U* (*n*) and matrix

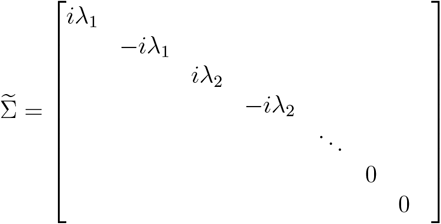

Satisfying 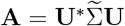. The 2*k*-th complex column vector **u**+*i***v** of **U** corresponds to the 2*k*-th and (2*k* +1)-th real column vectors **u** ∈ ℝ^*n*^ and **v** ∈ ℝ^*n*^ of the orthonormal matrix **P**. Geometrically, **A** acts on the real plane spanned by **u** and **v** via a *π/*2-radian (resp. —*π/*2-rotation) plus stretching by *λ*_*j*_for each pair of complex conjugate eigenvectors **u**(*j*) + *i***v**(*j*) (resp. **u**(*j*) — *i***v**(*j*)) in the columns of **U**.

##### Spectral theory and offset recovery

Suppose that a chain of offsets *γ*: *S*^1^ → ℝ^*n*^ comes from the ellipse model (12) with components

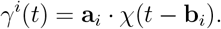

The difference of angles formula applied to the lead-lag matrix in 13 yields

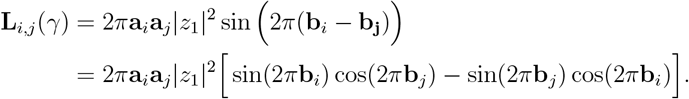

Denote amplitude-offset vectors by

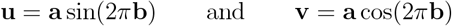

respectively. At the matrix level, the relation above implies that

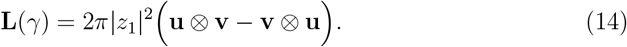

In this case, the lead-lag matrix is a rank 2 matrix, whose coimage (i.e., the span of its row vectors) is concentrated in the **uv**-plane. As per the discussion above, the fact that **L**(*γ*) is skew-symmetric implies that it must be a rotation in this plane. The eigenvalues corresponding to a non-zero complex-conjugate pair of eigenvectors can be calculated in terms of **u** and **v** as

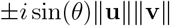

where *θ* is the angle between **u** and **v**. Following [16], the non-trivial complex eigenvector **z** = Re(**z**) + *i*Im(**z**) ∈ ℂ^*n*^ of **L**(*γ*) has components

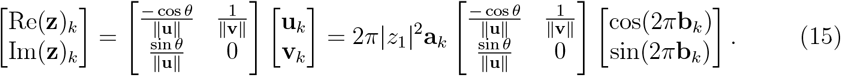

This formula implies the following important point, as noted in [16]:

*The amplitudes and offsets of dimensions in the ellipse model are related to the principal eigenvector of the lead-lag matrix by a linear transformation*.

Hence, one recovers the cyclic order of the coordinates.

###### Remark 12

*By 15, the extent to which the amplitude is recoverable depends on the angle θ between* **u** *and* **v**. *As a heuristic justification, we note that* sin *and* cos *are orthogonal as functions, with a further mathematical discussion of when the amplitude estimates are reliable to future work*.

#### Simplicial cyclicity analysis

We present here our generalization of cyclicity analysis to embedded simplicial complexes.

##### Manifold model

Let *M* be an *m*-dimensional oriented Riemannian manifold with volume form Vol_*M*_, and denote its Hodge star operator by

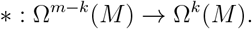

This induces an inner product on *k*-forms defined by

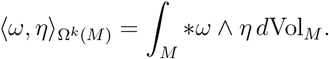

For a 1-form *η* ∈ Ω ^1^(*M*) and a smooth map *φ*: *M* → ℝ^*n*^, the *manifold lead-lag* is given by

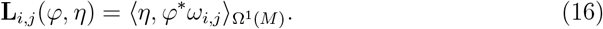

where

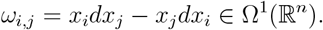

Path lead-lag is a special case of manifold lead-lag. Let *dθ* = ^*^1 be the volume form on *S*^1^ with the standard Riemannian structure, *γ*: *S*^1^ → ℝ^*n*^ a closed path, and **L**_*i,j*_(*γ*) the standard lead-lag matrix of the path *γ* given in Equation (8).

###### Proposition 13

**L**_*i,j*_(*γ*) = **L**_*i,j*_(*γ, dθ*).

*Proof*. We have

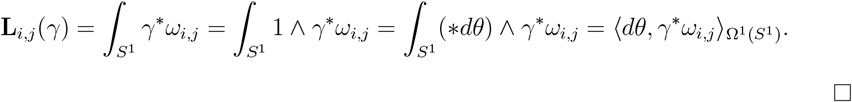

##### Simplicial model

Let *φ*: 𝒮 → ℝ^*n*^ be an oriented simplicial complex affinely embedded in ℝ^*n*^. For an edge [*x*_*p*_, *x*_*q*_] ∈ 𝒮, define

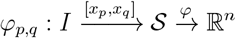

to be the path from *φ* (*x*_*p*_) to *φ* (*x*_*q*_). Integration induces a simplicial cochain

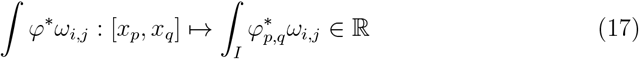

in *C*^1^(𝒮; ℝ) for each 1 ≤ *i, j* ≤ *n*. Equip *C*^1^(𝒮; ℝ) with the inner product 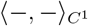 from Equation (4) which is induced by the geometry of the ambient ℝ^*n*^.

###### Definition 14

*The simplicial lead-lag matrix over α* ∈ *C*^1^(𝒮; ℝ) *is*

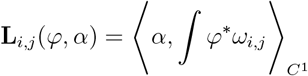

We reason that since 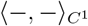 is an approximation of the underlying Riemannian metric (when it exists), the simplicial lead-lag is an approximation of the manifold model (when the data indeed lies on a manifold).

##### Geometric interpretation

For an affinely embedded simplicial complex, a simple application of Stokes’ theorem shows that the integrated cochain in Equation 17 has the following interpretation.

###### Proposition 15

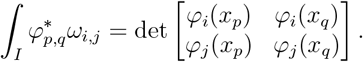

This corresponds to the signed area in the *ij*-plane of the parallelogram spanned by vectors *ϕ*(*x*_*p*_) and *ϕ*(*x*_*q*_). In practice, the simplicial complex will be the Rips complex *𝒳*_ε_of a normalized count matrix **X** conceptualized as a point cloud in ambient gene expression space. If we use the inner product on simplicial 1-cochains described by Equation 4, the coefficients of the lead-lag matrix are given by the following formula:

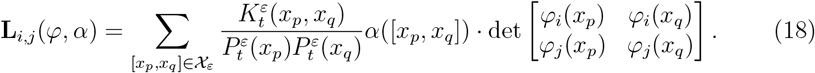

This quantity represents a type of *α-weighted signed area*. In other words, it is a linear combination of the signed areas in the *ij*-plane of parallelograms spanned by *φ* (*x*_*p*_) and *φ* (*x*_*q*_) for edges [*x*_*p*_, *x*_*q*_] in the Rips complex, weighted by the 1-cochain *α* and density specified by the inner product.

##### Change-of-basis

Let (*x*_1_,..., *x*_*n*_) be the standard coordinate functions on ℝ^*n*^, and

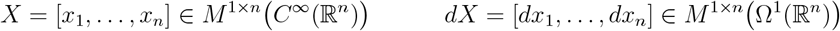

These matrices generate the lead-lag forms

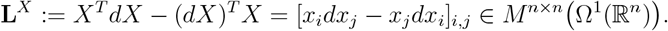

Suppose *y*_1_,..., *y*_*n*_is another orthornormal coordinate system on ℝ^*n*^ such that *Y* = *X***P** for some orthonormal matrix **P** ∈ O(*n*). Similarly, define the matrix of lead-lag forms with respect to the system *Y* as

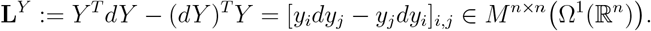

Let *φ*: 𝒮 → ℝ^*n*^ be an affine embedding and *α* ∈ *C*^1^(𝒮; ℝ) a simplicial 1-cochain. The lead-lag matrices with respect to the two coordinate systems are given by

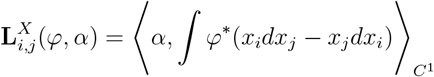

and

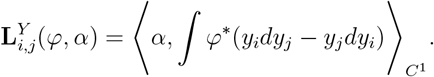

They are related by the following formula.

###### Proposition 16

**L**^*Y*^ (*φ, α*) = **P**^*T*^ **L**^*X*^(*φ, α*)**P**

*Proof*. We have

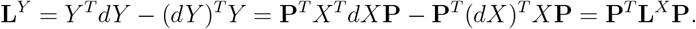

The linearity of integration, pullbacks and the operator 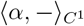 implies that

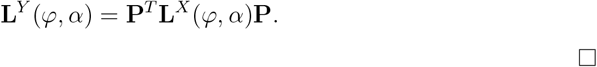

##### Spectral theory

Now suppose we have computed the lead-lag matrix **L**^*X*^(*φ, α*) with respect to an embedding, a 1-cochain, and the standard coordinates *X* on ℝ^*n*^. Putting our matrix in canonical form yields a real orthonormal matrix **P** ∈ *O*(*n*) and a block diagonal *n* × *n* matrix

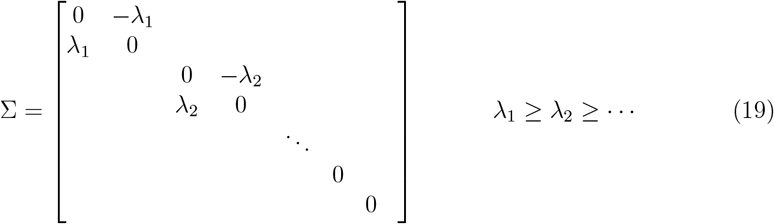

satisfying **L**^*X*^(*φ, α*) = **P**^*T*^ **∑ P**. Define a new coordinate system on ℝ^*n*^ by

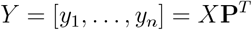

for *X* = [*x*_1_,…, *x*_*n*_] as above. A consequence of Proposition 16 is that we can interpret ∑ from Equation (19) as a lead-lag matrix

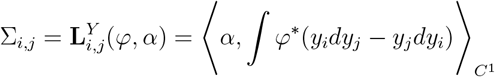

in the *Y* coordinates. In light of the discussion above of *α*-weighted signed area (18), we have the following interpretation:

*the value λ*_*i*_*in Equation* (19) *corresponds to the α-weighted signed area in the plane panned by y*_2*i*_*and y*_2*i*+1_.

Thus, putting our lead-lag matrix in canonical form identifies the plane spanned by *y*_0_and *y*_1_as the *principal plane*, i.e., that which maximizes the *α*-weighted signed area. Further, the ratio *λ*_1_*/λ*_2_describes the extent to which the *α*-weighted signed area is fully captured by the principal plane.

##### Recentering

A major difference between simplicial and path cyclicity analyses is that simplicial cyclicity analysis is not invariant under translations of the embedded simplicial complex. To mitigate the effect of translations, we first perform a procedure to “recenter” the data around the cochain *α*. Given a 1-cochain *α*, define a weighting on the vertices by 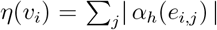 for the cochain *α*_*h*_evaluated on each oriented 1-simplex *e*_*i,j*_in 𝒮. The *α*-weighted mean is defined as

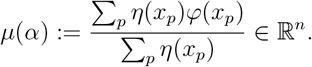

The recentered data is defined to be 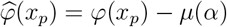 for all *p*.

## Discussion

In cases where data is sampled from *S*^1^, and *α* approximates the volume form, we claim that the simplicial lead-lag is a good model of the path lead-lag. In this case, it is reasonable to expect that the spectral decomposition of the simplicial lead-lag matrix exhibits a similar connection with the offsets in the Fourier decomposition of the chain-of-offsets model. In order to rigorously generalize this relationship, one needs a version of the chain-of-offsets model for manifolds rather than paths. We plan to define and study such a model in future work.

### Cyclicity analysis on single-cell data

The goal of this section is to contextualize Figure 2 in simplicial lead-lag theory.

#### Dominant geometric data

Recall that we denote the first *k* components of PCA projection of a single-cell experiment *X* calculated in the subspace spanned by genes 𝒢 by 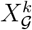. Our principle is that 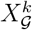 is the basic geometric model of the dominant biological phenomena in 𝒢. The points of the metric space 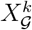 are in bijection with the points in *X* with the metric induced by the PCA projection 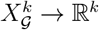 (Figure 2A).

The following geometric data are associated with this choice of metric.

1. The persistent cohomology 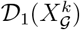 and ring score 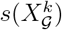
2. The dominant interval 𝕀_[ε,*δ*)_and associated Rips complex 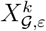
3. Inner-products over 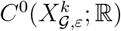 and 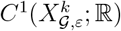, via Equation (4)
4. Harmonic representative 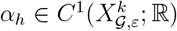 and circular coordinate 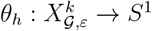

The harmonic representative *α*_*h*_is a simplicial model for the locally-defined gradient vector field^2^ of *θ*_*h*_in the Riemannian case (Figure 2B). A key assumption is that this is a model for the likely circular dynamics in the data, noting that orientation is arbitrary and needs to be determined by the user based on biological information.

#### Lifting

The geometric data model the underlying biological process in 𝒢. To understand which genes influence this process, we examine the lift (Figure 2B)

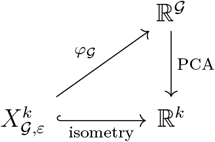

where *φ*_*G*_is map that linearly extends the expression of genes *G* at each node to the Rips complex at parameter *ε* ∈ ℝ. The lead-lag matrix (Figure 2C)

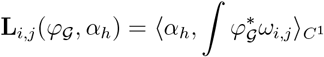

for 1 ≤ *i, j* ≤ |𝒢| represents pairwise gene ordering relative to *α*_*h*_. The lead-lag projection and principal complex eigenvector components approximate the global order in which the genes in 𝒢 occur along *α*_*h*_(Figure 2D).

## Data and code availability

The cHunter software package and the Jupyter Notebooks to analyze the publicly available data are available online at Github. The original human dermal fibroblast data will be deposited in GEO under accession GSE312110.

## Author contributions

K.M., M.K.Y. and K.H conceived the study. K.M and M.K.Y developed the overall analysis framework and led the project. K.M. conceived of topological and geometric framework, implemented the cHunter software, performed the topological and geometric analyses and generated the figures. M.K.Y. contributed to the development of the circular enrichment framework, implemented key algorithms and performed computational analyses. T.J.N. assisted with synthetic experiments and design. J.I. designed and carried out the experimental work. J.I, W.K. and C.P provided biological expertise and interpretation of results. M.A. contributed to data curation, software development and reproducible analysis workflows. H.A.H. and K.H. supervised the mathematical and computational aspects of the work. G.P.D. supervised the biological and translational aspects of the work. K.M., M.K.Y. and G.P.D. wrote the manuscript with input from all authors. All authors discussed the results and approved the final version of the manuscript.

## A. Experiment and pre-processing details

### A.1 Synthetic Transcriptomic Switch Model

#### Classical model

Our main synthetic model was a modified version of the linear ODE model of gene transcription and translation in [1]. The original linear model in [1] is governed by two ODEs:

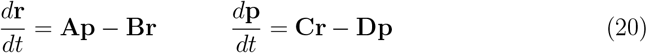

where

1. **r** and **p** are the vectors of mRNA and protein levels of the *n* genes at a given time.
2. The *n* × *n* interaction matrix **A** = [*a*_*i,j*_]_*i,j*_describes the linear effect of protein *j* on the RNA production of gene *i*.
3. The matrices **B, C** and **D** are diagonal matrices corresponding to degradation rates of RNA, and translation and degradation rates of proteins, respectively.

#### Transcriptional switch model

RNA transcription is known to occur in bursts [2, 3]. More recent models of RNA transcription account for this by designating discrete transcriptional states with constant transcription rates [4–7]. To adapt the linear ODE model in (20) to this phenomenon, as well as to constrain the RNA counts to finite quantities, we introduced a threshold function

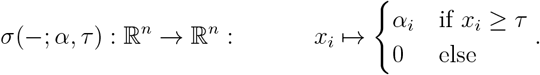

The parameters *α*_*i*_correspond to transcriptional rates for gene *i* given that the interaction is above certain threshold *τ* ∈ ℝ. Our updated transcriptional switch model is then given by

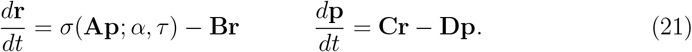

Given initial conditions **r**_0_and **p**_0_, solutions for the above system of ODEs were approximated using the `scipy.integrate` package over a finite sequence of discrete times

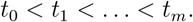

Example solutions to this model are given in Figure S1D and Figure S2D.

#### Negative feedback loops

We first wanted to connect theoretical ODE models of RNA transcription to the emergence of global topological circular structures. Based on the chain-of-offsets model in (see Cyclicity Analysis in the Method’s section), we hypothesized that when the protein-RNA interaction matrix **A** in the transcriptomic switch model 21 is a skew-symmetric rank 2

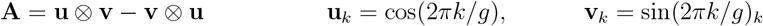

There would be an evident circular feature in low-dimensional PCA. Our rationale was that such processes model a dominant negative feedback loop in the data whose dynamics is described by the principal complex eigenvector of **A**. Despite the modified transcriptional switch model and realistic noise, the lead-lag matrix (Figure S2B) provided a relatively robust model of the relationships between different genes (Figure S2C).

#### Transient processes

We also wanted to demonstrate the viability of using circular enrichment in studying transient processes that are not necessarily periodic. To recreate such a process in our model, we designed an interaction matrix where the earlier genes positively influenced later genes and vice versa (Figure S1). Circular gene set enrichment revealed significant circular structure outside of the first principal component (Figure S1G,H). The reconstructed phase estimates reliably recovered the ordering of genes (Figure S1I,J), and the lead-lag matrix provided a reasonable estimation of the underlying gene-protein interaction matrix (Figure S1C).

#### Model parameters

In Table 8 we present the parameters for the transient (T) and recurrent (R) examples (Figure S1 and Figure S2) of the synthetic transcriptomic switch model. Parameters were manually selected to lead to approximate ODE solutions that were sufficiently rich. Sensitivity analysis (Figure S2D) parameters (S) of the recurrent example were adjusted as a function of the number of cells and genes. Ground-truth cell times were selected by uniformly dividing the time interval, so that the gene expression of a cell at time *t* corresponds to the ODE solution at time *t*.

**Table 8:** Hyperparameters in the transcriptomic switch model experiments.

| | $g$ | $c$ | $t$ | $\mathbf{r}(0)$ | $\mathbf{p}(0)$ | $\tau$ | $\alpha$ | $\mathbf{B}$ | $\mathbf{C}$ | $\mathbf{D}$ |
| --- | --- | --- | --- | --- | --- | --- | --- | --- | --- | --- |
| T | 80 | 1000 | $[0, 6)$ | $\begin{cases} 5 & i \leq 10 \\ 0 & \text{else} \end{cases}$ | 0.0 | 10.0 | $\sim U(1, 4)$ | $\sim U(1, 1.6)$ | $2 \cdot I_{80}$ | $I_{80}$ |
| R | 40 | 1000 | $[0, 20)$ | $\begin{cases} 5 & i \leq 10 \\ 0 & \text{else} \end{cases}$ | 0.0 | 3.0 | $\sim U(1, 4)$ | $\sim U(1, 1.6)$ | $2 \cdot I_{40}$ | $I_{40}$ |
| S | $g$ | $c$ | $[0, 0.02c)$ | $\begin{cases} 5 & i \leq 0.1g \\ 0 & \text{else} \end{cases}$ | 0.0 | $0.02g$ | $\sim U(1, 4)$ | $\sim U(1, 1.6)$ | $2 \cdot I_g$ | $I_g$ |

#### Sampling with negative binomial noise

To realistically model noisy and stochastic single-cell data, we adopted the approach of [8] of negative binomial sampling, which has been observed to be a good model for the distribution of RNA molecules [2, 3]. Our approximate solutions of mRNA levels according to Table 8 were used as the mean in a negative binomial sampling with a constant dispersion parameter to generate a random count matrix that converges to our solved system in log mean (Appendix A.1). Recall that the negative binomial distribution NegBinom(*n, p*) has probability mass function

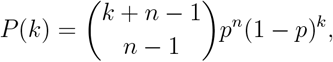

where *k* is the number of failures over *n* trials with probability *p* of success. The parameters are related to the mean *µ* and variance *σ*^2^ by the formulae

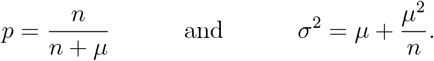

The approximated solutions of mRNA counts (**r**_*i*_(*t*_*j*_)) in Table 8 over discrete time intervals *t*_0_*< t*_1_*<*…*< t*_*n*_were exponentiated, and used as the mean in a negative binomial sampling with a constant dispersion parameter *d* = 1*/n*. This generated a random count matrix

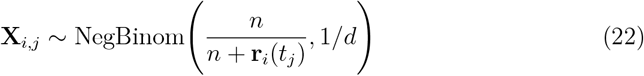

that converges to our solved system in the mean. Data were sampled from this using the Python function `scipy.stats.nbinom.rvs`. Examples of the sampling are given in Figure S1E-G and Figure S2E-G.

### A.2 Benchmarking

#### Velocycle synthetic model

We benchmarked the estimation of circular coordinates against the synthetic model presented in [8]. In this model, mean spliced RNA expression is modeled directly as the first harmonics in the Fourier series:

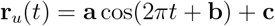

where 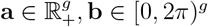 and **c** ∈ ℝ^*g*^ are the respective vectors of amplitudes, phases and offsets of each gene. Each of the parameters **a, b** and **c** are drawn from a normal distribution with predefined mean and correlation structure (see [8] for details), and each cell *i* is associated with a pseudotime *t*_*i*_, drawn from a uniform distribution *U* (0, 1). As for the previous examples, the vector **r**(*t*) is used as the mean in the negative binomial distribution for sampling the data as per Equation (22), but this time with a stochastic dispersion parameter given by

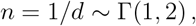

where Γ(*α, β*) is the gamma distribution with shape parameter *α* and rate parameter *β*.

#### Velocycle benchmarking

The pipeline for our model in the `velocycle` benchmarking (Figure S3A and C) consisted of: separating each gene into spliced and unspliced raw UMI counts, filtering genes that occurred in fewer than three cells, normalizing the per-cell raw UMI counts, and projecting onto the first two PCA coordinates, followed by computing the persistent cohomology circular coordinate of the most persistent feature. The PCA baseline consisted of identical preprocessing, with the circular coordinate taken as the angle between the first two principal components. For each experiment, we ran `velocycle` with its suggested preprocessing and parameters for 200 epochs [8]. The box-plots in Figure S3C were run with the same parameters 20 times with 3000 cells and 300 genes, as per [8].

#### Multi-process benchmarking

The multi-process benchmark was built from a minor adaptation of the `velocycle` synthetic data generation framework. This benchmark uses a dataset containing two independent processes (linear and circular) to evaluate the performance of `cHunter`, `PCA` and `velocycle` with an incorrect prior, i.e., the underlying manifold assumed is a circle, *S*^1^, whereas by construction this dataset has the shape of a cylinder, *S*^1^ × *I*.

For a fixed number of cells *c* and genes *g*, we generated the standard `velocycle` dataset to attain a circular process *X*_circ_⊂ ℝ^*c*×*g*^. We then generated another, independent, `velocycle` dataset with 2*c* cells and *g* genes, before restricting to the first half *c* cells ordered by phase to create a non-circular process modelled by a half-circle *X*_arc_⊂ ℝ^*c*×*g*^. The two datasets were then shuffled and concatenated along the cell axis, to create a dataset consisting of separate uncoupled circular and linear processes with *c* cells and 2*g* genes *X*_total_⊂ ℝ^*c*×2*g*^. All `cHunter` estimates were carried out in the concatenated unspliced-spliced phase space.

### A.3 Cycling dermal fibroblasts

#### Dataset

The data we used from [9] is publicly available on the Gene Expression Omnibus database under accession no. GSE167209. We followed the pre-processing procedure in [8], where the same dataset is used as an example. Similar to our pipeline and the standard Seurat pipeline, this consisted of log normalization and mean centering, however, with additional filtering for both spliced and unspliced genes.

### A.4 AR silencing in dermal fibroblasts

#### Dataset

Our single-cell RNA-seq dataset consisted of human dermal fibroblasts (HDFs) infected with two lentiviruses expressing different AR-silencing shRNAs versus a control expressing scrambled shRNAs. Following puromycin selection, cells were collected and processed for single-cell RNA-seq using the 10x Chromium 3’ kit according to the manufacturer’s instructions. Sequencing was subsequently performed at the Lausanne Genomic Technologies Facility.

#### Preprocessing

Pre-processing and filtering were performed by grouping the shAR conditions with the control. This dataset consisted of 5,607, 3,663, and 4,410 cells in the shAR1, shAR2, and CTRL subgroups, respectively. We first performed clustering via the Leiden algorithm and filtered three populations of cells based on abnormally low total RNA counts, mitochondrial percentage reads, and MALAT1 expression (Figure S5A,B,C). The rationale for grouping the conditions together was to (1) identify shared circular structures before (2) locating perturbations or deviations specific to each condition. In the combined dataset, cells expressing at least 200 genes and genes expressed in at least three cells were retained. Raw UMI counts were normalized per cell and log-transformed. Highly variable genes were then identified using `scanpy.tl.highly-variable`. For comparison, we used scanpy’s default cell-cycle scoring to assign G1, S and G2M labels to each cell (Figure S5D,E,F). Notably, while this overall cell cycle scoring showed similar G1, S, and G2/M distributions across the bulk populations of control and AR-silenced cells, further analysis would reveal specific cell-state changes linked to AR silencing (see below).

### A.5 Prostate regeneration data

#### Dataset

Original data profiles were from 13,398 cells of the mouse prostate, focusing primarily on the anterior lobe, using droplet-based scRNA-seq. For recovering cell type classifications, we used the author’s data and annotations for [10] available at Gene Expression Omnibus https://www.ncbi.nlm.nih.gov/geo/ (accession no. GSE146811). We restricted the data to the largest cluster of luminal epithelial cells (L1), characterized by the genes *CD24a, Krt8*, and *Krt18*.

#### Pre-processing and filtering

We first restricted to the subset of the Luminal 1 (L1) subpopulation, then filtered by cells expressing at least 200 genes and genes expressed in at least three cells. Raw UMI counts were normalized per cell, log-transformed, and restricted to the subset of highly variable genes using `scanpy.tl.highly-variable`. MAGIC [11] was applied to the data with nearest-neighbour parameter knn = 4 for one step (*t* = 1). Finally, we removed 3000 outliers with low density, according to the density function

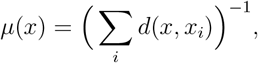

where *d* was the metric in the first five principal components. Finally, we performed maxmin sampling of the data to reduce it to a set of 2^10^ landmark cells.

#### Circular enrichment

For the circular gene set enrichment, we used the 2020 hallmark gene set from the Molecular Signatures Database [12]. The global pre-processing was first applied to the luminal cells above. We then projected onto each gene set, followed by the first two principal components, applied iterative density filtering and finally computed the circular coordinate of the most persistent feature. The *p*-values were computed according to the permutation procedure described in Method.

## Footnotes

1 Magic is a low-pass filter which replaces the gene expression of each cell with a weighted average of its nearest neighbours.

2 Formally, the dual vector field of the associated 1-form.

