## Supplementary figures for "Topology identifies concurrent cyclic processes in single-cell transcriptomics and androgen receptor function"

### B Supplementary Figures

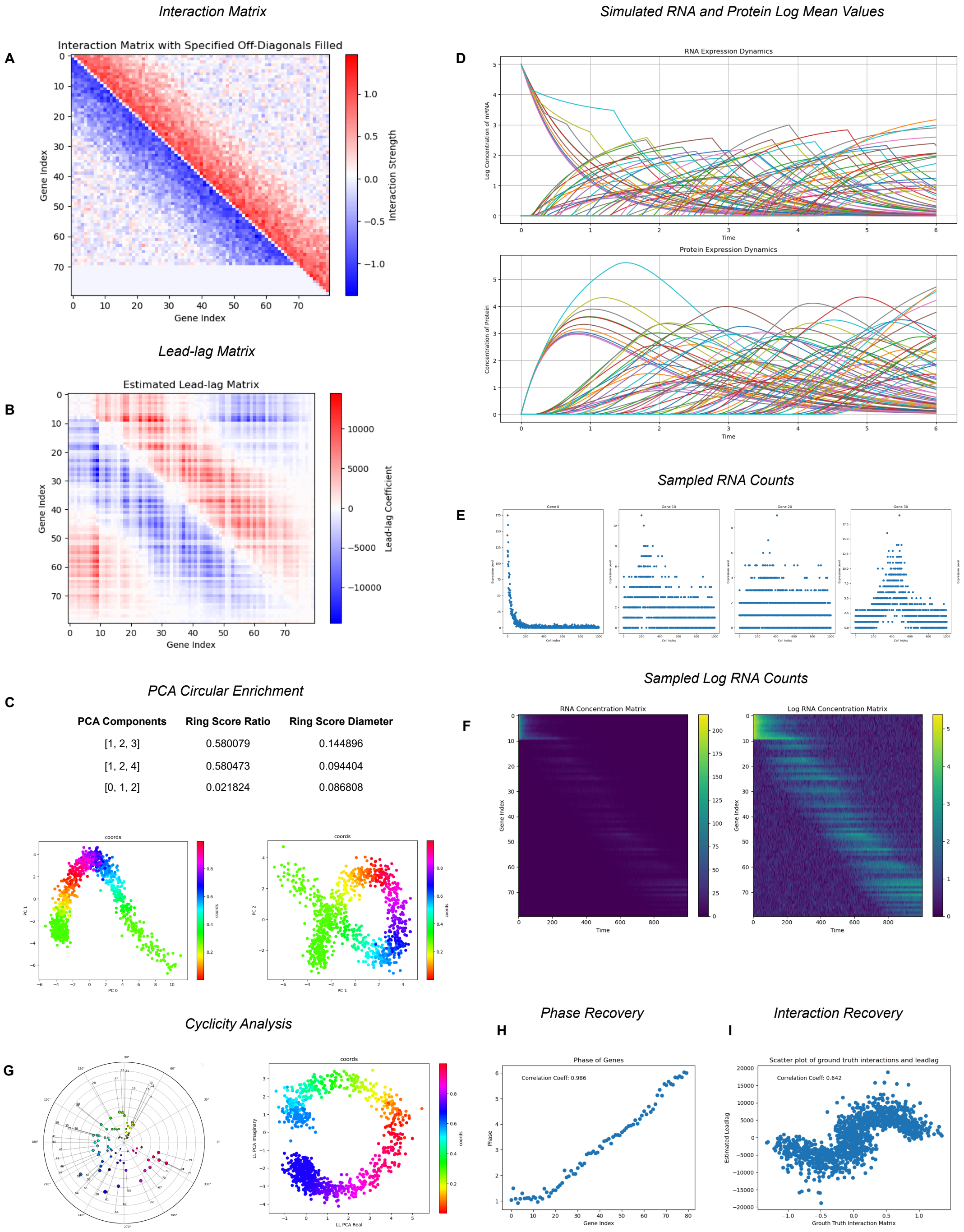


Figure S1: Analysis of a synthetic, transient mRNA transcription event. **(A)** The synthetic interaction matrix **A** of a transient mRNA transcription event. **(B)** The estimated lead-lag matrix. **(C)** PCA circular enrichment and selected projections onto principal components. **(D)** The simulated log means of RNA and protein. **(E,F)** The negative binomial sampled RNA counts and their log transforms. **(G)** Cyclicity analysis-based phase estimation and lead-lag projection. **(H)** The estimated gene phases from cyclicity analysis and their correlation with the ground-truth gene ordering. **(I)** The values of the synthetic interaction matrix plotted against their values in the lead-lag matrix.

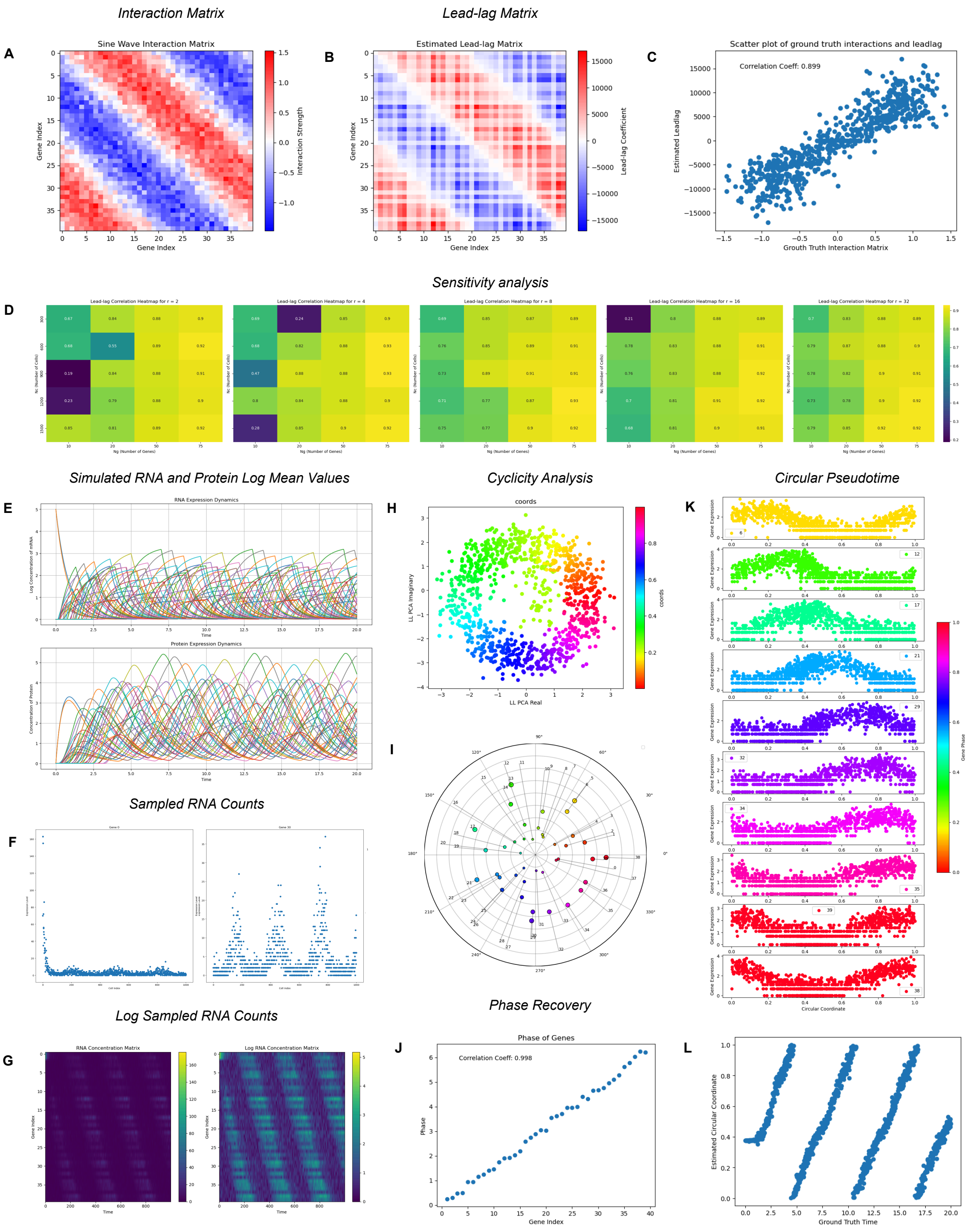


Figure S2: **(A)** The synthetic interaction matrix **A** of a transient mRNA transcription event. **(B)** The estimated lead-lag matrix. **(C)** The values of the synthetic interaction matrix plotted against their values in the lead-lag matrix. **(D)** Sensitivity analysis over number of cells, genes and dispersion parameter. **(E)** The simulated means of RNA and protein. **(F,G)** The negative binomial sampled RNA counts and their log transforms. **(H-I)** Lead-lag projections and cyclicity analysis estimated gene phases. **(J)** Correlation of estimated gene phases with ground-truth gene orderings. **(K)** Log expression over estimated circular-pseudotime. **(L)** Ground-truth time against estimated circular coordinate.

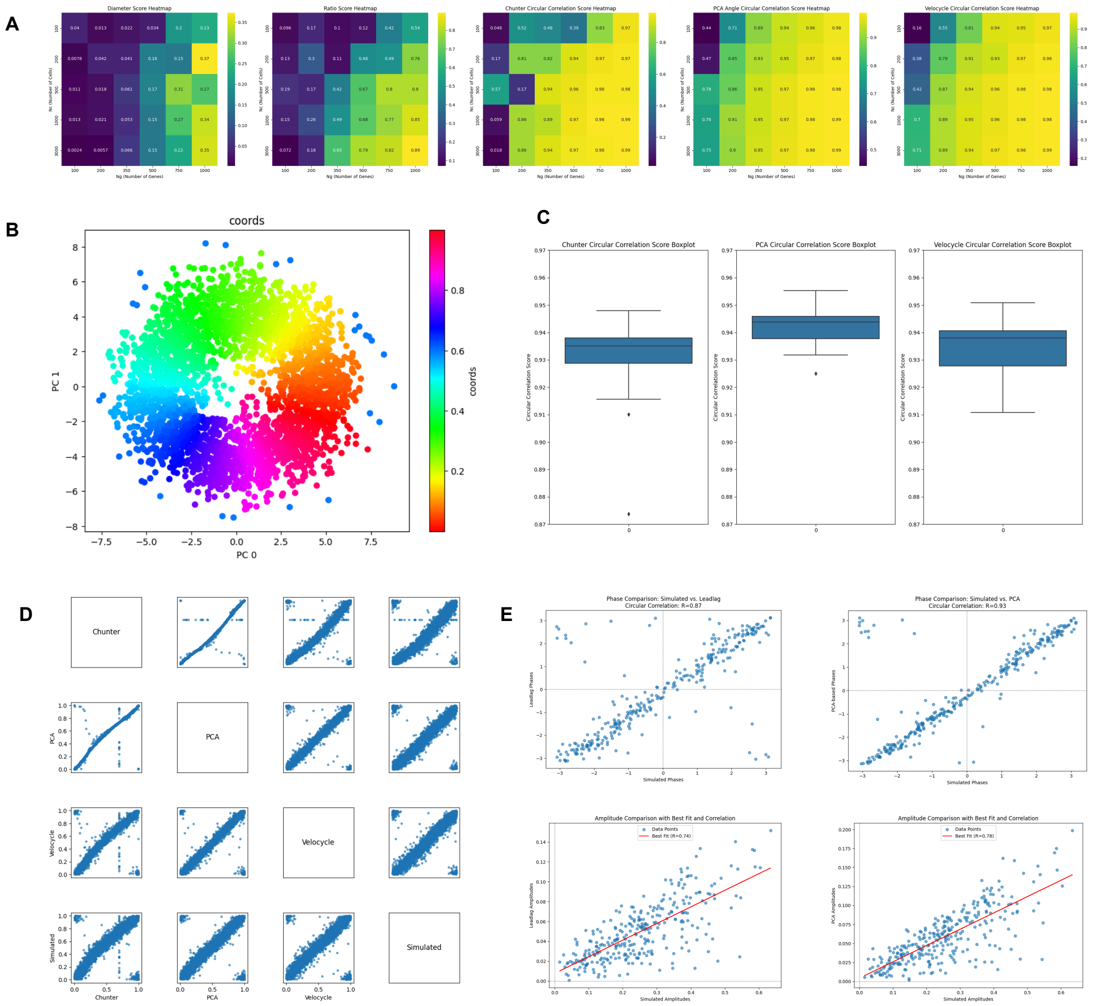


Figure S3: **(A)** Left: diameter and ratio ring scores for PCA projection of synthetic model. Right: circular correlations with ground-truth circular coordinates using different methods. **(B-E)** Case study of synthetic model with 3000 cells and 300 genes. **(B)** PCA projection in first two components **(C)** Circular correlation with ground-truth box-plot over 20 runs using cHunter, PCA baseline and Velocycle. **(D)** Pairwise comparison of different circular coordinate estimates. **(E)** Ground-truth genewise Fourier phases (top) and amplitudes (bottom) vs estimates in cHunter and PCA baseline.

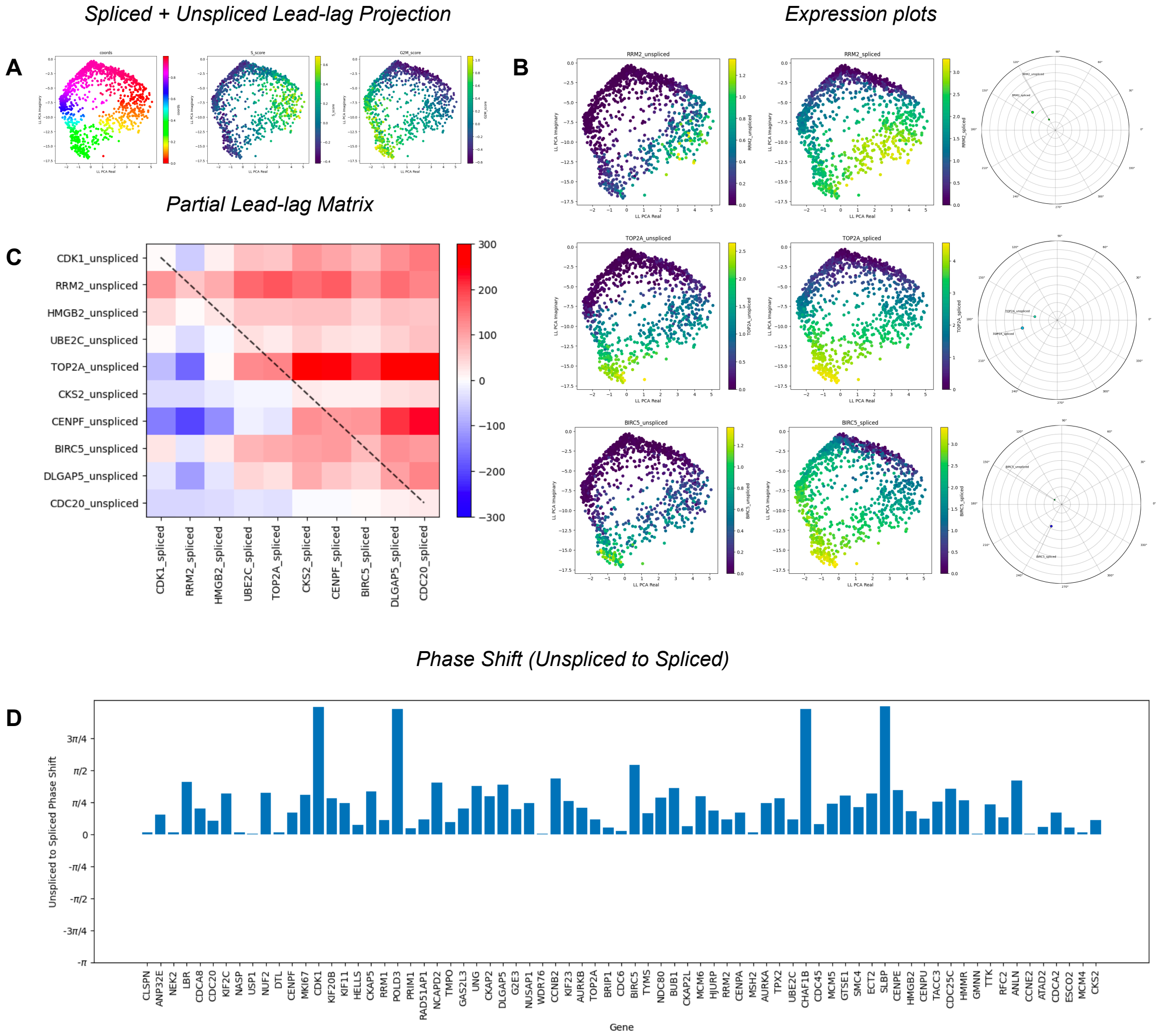


Figure S4: **(A)** Lead-lag projection of the spliced and unspliced cell-cycle gene set expression data using the principal circular feature. **(B)** The spliced and unspliced expression plots of RRM2, TOP2A and BIRC5 and their estimated phases from harmonic lead-lag analysis. **(C)** The lead-lag matrix of the top 10 genes, displaying only the lead-lag of unspliced against spliced. **(D)** The phase shift from the estimated phase of unspliced to spliced of each gene in the cell cycle gene set.

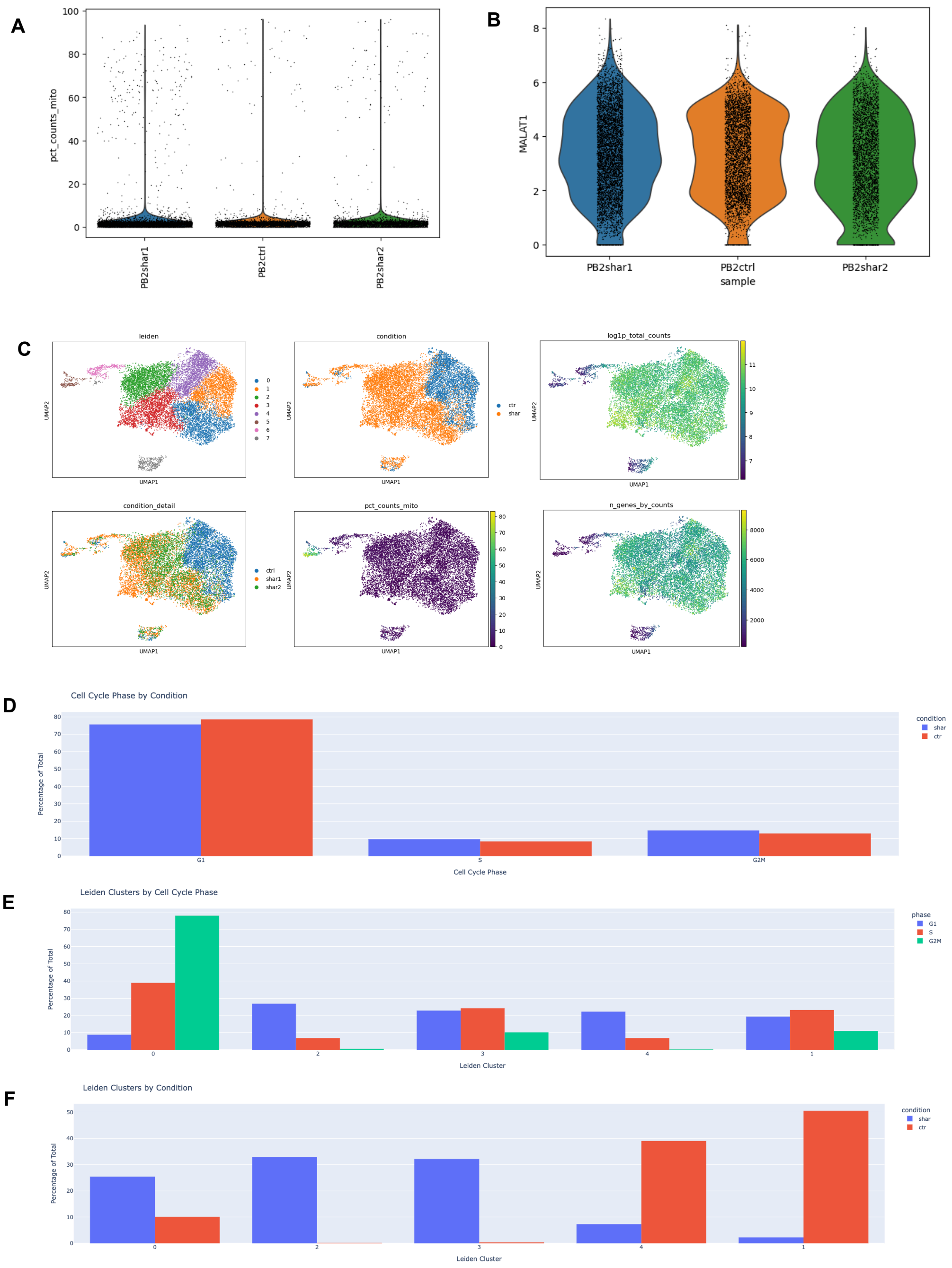


Figure S5: Pre-processing and clustering of AR-silenced human dermal fibroblasts. **(A)** Violin plots of the percentage of non-zero genes mitochondrial across conditions. **(B)** Violin plots MALAT1 across conditions for quality control. **(C)** UMAP plots colored by Leiden clusters, conditions and various quality control metrics. **(D)** Distribution of `scanpy`-derived cell cycle assignments across conditions. **(E)** Distribution of `scanpy`-cell cycle phase estimates across Leiden clusters. **(F)** Distribution of experimental conditions across Leiden clusters.

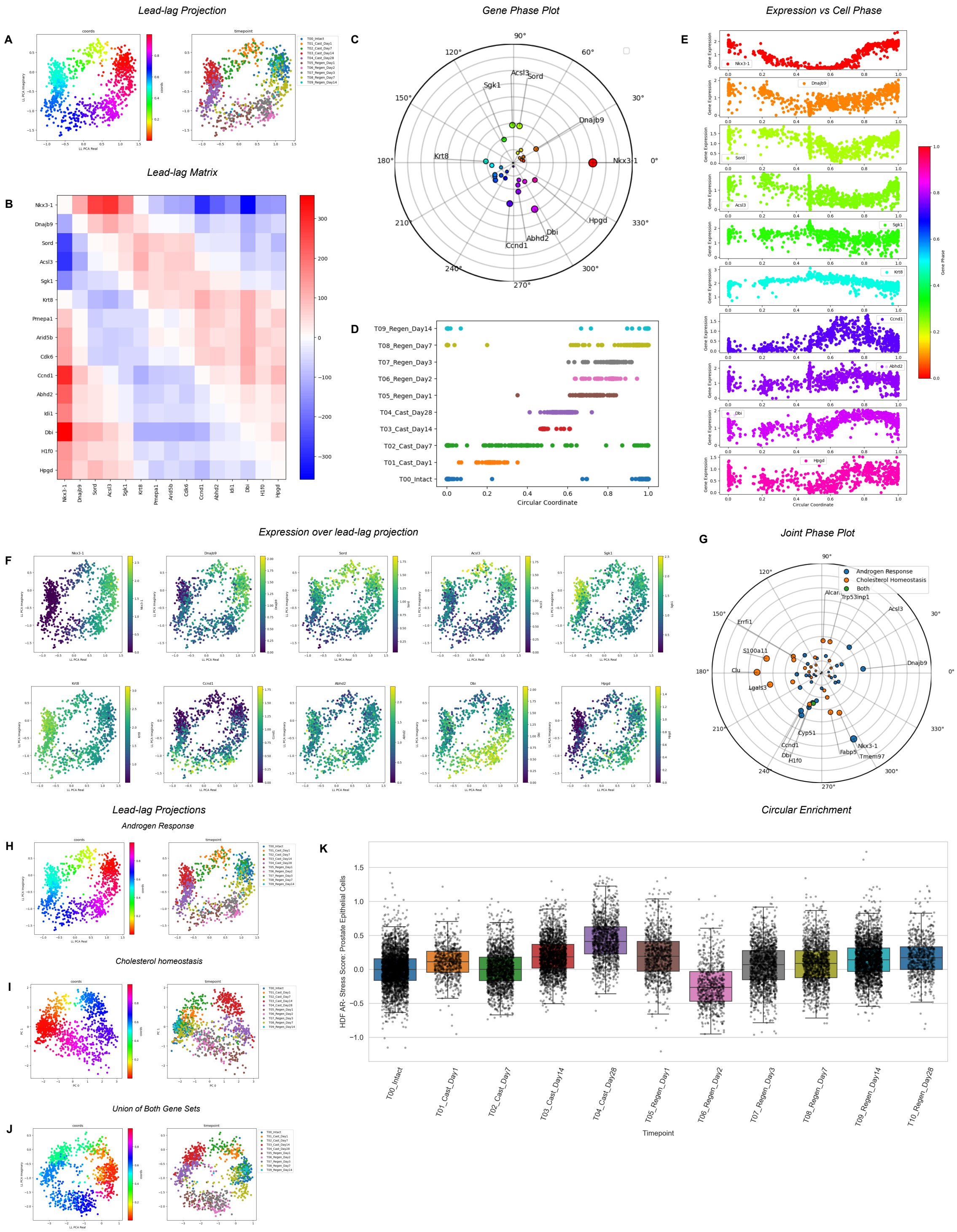


Figure S6: Analysis of prostate regeneration data. **(A)** The lead-lag projection colored by circular coordinates and sample time, respectively. **(B)** Lead-lag matrix. **(C)** The estimated gene phases. **(D)** Histogram plots of sample times distributed against the circular coordinates.**(E)** Gene expression of primary genes vs circular coordinate. **(F)** Gene expression of primary genes vs lead-lag projection. **(G)** Phase plot of genes against the circular coordinate attained in the union of the two gene sets. **(H-J)** The lead-lag projection against two enriched gene sets and their union, and corresponding ring score permutation tests and persistence diagrams. **(K)** Box plot of AR stress score against ground-truth time-points.
